# Weak selection and stochastic processes limit the emergence of antigenic variants during household transmission of influenza A viruses

**DOI:** 10.1101/2025.11.04.686470

**Authors:** Hunter J. Ries, Joseph Lalli, Kelsey R. Florek, Shari Barlow, Maureen Goss, Richard Griesser, Tonya Danz, Amra Uzicanin, Jonathan Temte, Thomas C. Friedrich

**Affiliations:** Department of Pathobiological Sciences, University of Wisconsin–Madison, Madison, Wisconsin, USA; Wisconsin State Laboratory of Hygiene, Madison, Wisconsin, USA; Department of Family Medicine and Community Health, University of Wisconsin–Madison, Madison, Wisconsin, USA; National Center for Emerging and Zoonotic Infectious Diseases, US Centers for Disease Control and Prevention, Atlanta, Georgia, USA (retired); Wisconsin National Primate Research Center, University of Wisconsin–Madison, Madison, Wisconsin, USA

**Author notes:** Correspondence should be addressed to T.C.F.

**Keywords:** Influenza A virus, Antigenic drift, Hemagglutinin antigenic sites, Transmission bottleneck, Intra-host single-nucleotide variants, Purifying selection, Household transmission, Whole-genome sequencing, A(H1N1)pdm09, A(H3N2)

## Abstract

Influenza viruses undergo antigenic drift, the gradual accumulation of mutations that cause antigenic changes in the viral surface proteins hemagglutinin (HA) and neuraminidase (NA). Although selection for antigenic variants is detectable on the global scale, the processes by which antigenic variants are generated and selected in individual hosts remain unclear. It has been hypothesized that selection for antigenic variants may occur during the establishment of a new infection, rather than over time in a single host. Here, we leveraged a large household cohort study to assess whether selection was detectable between acutely infected hosts. We investigated influenza A virus evolution using specimens from 384 children and household contacts with RT-PCR-confirmed influenza A infection, representing infections with A(H1N1)pdm09 and A(H3N2) viruses from 2017–19. In agreement with prior studies, we found that acute infections involved weak purifying selection across the viral genome. In addition, we identified 40 transmission events occurring in 31 households. During transmission, evolution between hosts was characterized by tight transmission bottlenecks and weak purifying selection. We found variability in the strength and direction of selection on antigenic regions of HA, but no clear evidence for selection of antigenic variants during transmission. Together, our results indicate that stochastic processes and weak natural selection dominate most acute influenza A virus infections and transmission events, and that selection of antigenic variants during transmission between acutely infected hosts is likely to be exceedingly rare.

**Author Summary:** Influenza viruses clearly evolve under selective pressure from immune responses in human populations, but recent work suggests that within individual infections random effects are stronger than selection. New viral variants that spread globally must nonetheless emerge in one person and be transmitted onwards—how does this happen? We characterized viral genomes collected over two influenza seasons from 384 children and their household contacts. We detected 40 transmissions among 31 of the households, allowing us to examine how selection acts during infection and transmission. We found that influenza virus genetic diversity is low in infected individuals, and mutations arising in one person are rarely transmitted to their household contacts, consistent with prior reports that influenza virus evolution is tightly constrained within hosts. We further examined all transmission events for evidence of selection between hosts, finding only one mutation that could plausibly affect antibody recognition. However, we found no evidence that this mutation was favored by natural selection. Our results suggest that chance events, together with weak selection, are the main forces affecting influenza virus evolution within and between hosts during typical acute infections. Selection for new variants may be more likely to occur over longer transmission chains and/or during prolonged infections.

## Introduction

Like other rapidly evolving RNA viruses, influenza A virus causes significant global morbidity and mortality [1]. Influenza vaccine effectiveness is often reduced by antigenic drift, the emergence and preferential propagation of antibody-escape mutations in the antigenic regions of the hemagglutinin (HA) and neuraminidase (NA) proteins. Influenza vaccine antigens must be frequently updated to reflect these changes in the HA protein of circulating influenza strains.

The National Institute for Allergy and Infectious Diseases has thus identified “accurately predicting how influenza viruses will evolve” as critical to developing an effective universal influenza vaccine [2]. Although influenza A virus evolution is detectable on the global scale, it remains unclear how new antigenic variants arise in one host and are propagated through transmission chains and among populations. An improved understanding of these processes will enhance our ability to forecast trends in viral evolution.

Variant viruses encoding one or more amino acid substitutions quickly arise within hosts after infection [3–7]. These intra-host single-nucleotide variants (iSNVs) cause nonsynonymous (amino-acid changing) or synonymous (“silent”) mutations, which may impact pathogenesis [8]. Purifying selection purges deleterious mutations from viral populations, while diversifying selection favors changes away from the within-host consensus sequence. Globally, seasonal influenza viruses show an excess of synonymous variants throughout most of the genome, suggesting that most nonsynonymous mutations are deleterious and are quickly purged from circulation [8]. Classically defined antigenic regions of HA are the exception and are characterized by an excess of nonsynonymous substitutions. This excess suggests that HA sequences are subject to positive selection at a global scale, which is reflected in the “ladder-like” HA phylogeny and the continual emergence of new HA clades [8]. However, the action of natural selection appears to be relatively weak within individual hosts, even in HA antigenic sites [8]. Studies by our group [3,4] and others [7–14] have thus far failed to detect evidence of selection for antigenic variants during acute infections, where even potential immune-escape mutations remain at low frequency. Interestingly, influenza viruses appear capable of accumulating antigenic changes during chronic infections in immunocompromised individuals [8,15,16], suggesting that acute infection may not allow enough time for antigenic variants to emerge [14].

Every positively selected variant must originate from a mutation in a single host [17,18]. What evolutionary processes allow some mutations to “escape” from the forces of randomness and weak selection in individual hosts to become dominant in the global population? Recently, Morris *et al.* [19] suggested that immune selection is strongest at the time a new infection is being established at a respiratory site in the presence of pre-existing virus-specific antibodies. In this model, incoming viruses are subject to “sieving” by pre-existing antibodies in the recipient, such that viruses capable of evading antibody detection are more likely to successfully found a new infection. Antigenic variants that are present at low frequency in a donor may therefore become fixed in a new host during a selective transmission bottleneck. In contrast, because antibodies specific for the incoming virus become detectable only near the end of a typical acute infection, there is a temporal disconnect between the peak of virus replication and the appearance of either recall or de-novo antibody responses capable of selecting for new antigenic variants. Transmission of these variants is further impacted by the transmission bottleneck, which involves a sharp reduction in viral diversity—according to most estimates, only 1–2 unique viral genomes from the donor establish infection in the recipient [7,20]. In their model, Morris *et al.* suggest that minor iSNVs under positive selection could reach 100% within-host frequency within the first 24 hours of infection [19]. In support of this, we have previously shown in a ferret model system that influenza A virus variants present at less than 10% frequency in the donor can survive the transmission bottleneck and reach fixation in a new host [21].

Detecting potential selection on viral genomes during the establishment of a new infection requires characterizing virus populations in donor-recipient pairs and cannot be done through studies of individual infections alone. Our objective was to determine whether immune selection for antigenic variants is detectable during acute, person-to-person transmission of influenza A virus. We hypothesized that if antibody-mediated sieving acts at transmission, it might be possible to detect events in which low-frequency antigenic variants in donors would become enriched or fixed in recipients. Here, we sequenced seasonal influenza A viruses from 384 influenza-positive individuals enrolled in a school-associated, community-based influenza surveillance study, the Oregon Child Absenteeism Due to Respiratory Disease Study (ORCHARDS; Oregon, Wisconsin, USA) [22,23]. We then quantified within-host diversity and selection, linked donors and recipients using epidemiologic and genetic criteria, and tested whether variants—particularly in HA antigenic sites—were preferentially transmitted to recipients.

## Results

### Study population and viral genome sequencing

For this study, we used specimens collected from participants in the Oregon Child Absenteeism Due to Respiratory Disease Study (ORCHARDS) study. Full methodology was reported previously [22,23], and the protocol was approved by the University of Wisconsin–Madison Health Sciences Institutional Review Board, with written consent from adults and parental consent plus assent for minors. Briefly, the study enrolled school-aged children from the Oregon School District (Oregon, Wisconsin, USA) who reported at least two acute respiratory symptoms (e.g., cough, sore throat, nasal congestion, runny nose, sneezing, or fever). Study staff performed home visits, collected specimens from the children, and offered to enroll other household members in a sub-study assessing within-household influenza transmission. Study staff then collected nasal swabs from any other participating household members. Respiratory specimens that tested RT-PCR-positive for influenza A virus were used for the work described here. In total, we sequenced 384 samples from the 2017–18 and 2018–19 influenza seasons, generating 283 complete viral genomes that passed our quality-control thresholds (see Methods) [**Table 1**]. The average Ct value for all samples included in our study was 27.2, with a standard deviation of 3.9, and a range of 18.52–35.37.

**Table 1.** Study demographics.

|  | 2017–18 |  | 2018–19 |  | 2017–19 |
| --- | --- | --- | --- | --- | --- |
|  | A(H1N1)<br>pdm09 | A(H3N2) | A(H1N1)<br>pdm09 | A(H3N2) | Total |
| Samples sequenced | 27 | 163 | 79 | 115 | 384 |
| Passing quality control<sup>1</sup> | 24 (89%) | 119 (73%) | 70 (89%) | 70 (61%) | 283 (74%) |
| Female | 11 (46%) | 55 (47%) | 34 (49%) | 39 (56%) | 139 (49%) |
| Under 18 years of age | 19 (79%) | 102 (86%) | 54 (77%) | 65 (93%) | 240 (85%) |
| Recent flu vaccination<sup>2</sup> | 13 (54%) | 47 (39%) | 46 (66%) | 47 (67%) | 153 (54%) |
| Ct value (+/- SD) | 27.2 (3.6) | 28.4 (4.1) | 26.4 (3.6) | 26.2 (3.6) | 27.2 (3.9) |
| Households with ≥2 flu-<br>positive members<sup>3</sup> | 3 | 16 | 12 | 5 | 36 |
| Samples from households<br>passing genetic criterion<sup>4</sup> | 6 | 28 | 27 | 10 | 71 |
| Two individuals | 1 | 8 | 10 | 5 | 24 |
| Three individuals | 0 | 4 | 1 | 0 | 5 |
| Four individuals | 1 | 0 | 1 | 0 | 2 |

<sup>1</sup>All segments mapped; ≥99% genome coverage and ≥100× depth (see Methods).
<sup>2</sup>Current-season influenza vaccination at specimen collection, confirmed in the Wisconsin Immunization Registry [23].
<sup>3</sup>Households with at least two influenza-positive members, based on epidemiologic data alone. These 36 households contributed 46 potential transmission pairs (36 donors, 82 individuals).
<sup>4</sup>Samples from households meeting both epidemiologic and genetic criteria for transmission. These 71 samples represent 40 transmission pairs involving 31 unique donors.

We first used deep sequencing to assess viral diversity within all samples. We mapped sequence reads using each season’s A(H1N1)pdm09 or A(H3N2) vaccine strain as a reference and identified intra-host single-nucleotide variants (iSNVs) in each sample using conservative frequency thresholds (3%) and quality control criteria (see Methods). Then, we aligned all high-quality participant consensus sequences for each season and subtype and used the majority nucleotide at each position in this alignment to define a “within-season” reference sequence. Raw sequencing reads were then re-processed and aligned against the within-season reference sequence to contextualize the variants relative to the strains in local circulation at the time of sampling. We included the major genes in the 6 largest viral gene segments in our alignment and variant-calling pipeline. We excluded M1/M2 and NS1/NEP genes because their overlapping reading frames can confound site-based diversity metrics [8]. Some samples contained very high numbers of iSNVs, which could reflect truly high levels of within-host diversity, but may also indicate co-infection with distinct strains or possibly method errors. To remove such outliers, as in [8], we quantified the number of iSNVs (>0.5% allele frequency) for each sample, ranked them in descending order within their season-subtype group, and then excluded samples that were in the top 10%.

### Low-frequency variation and purifying selection dominate within-host viral evolution

To assess within-host diversity in our cohort, we plotted minority iSNV frequencies (3–50%) for each gene and annotated them by mutation type: nonsynonymous (amino-acid changing), synonymous (“silent”), or stop gained (**Fig 1A**). We detected nonsynonymous and synonymous iSNVs across the genome, with a large proportion (77.3%) of iSNVs detected below 10% allele frequency in all samples. The pattern of mostly low-frequency iSNVs found throughout the viral genome was consistent across influenza seasons and viral subtypes (**S1 Fig**). We then quantified the total number of iSNVs per sample: 68 samples contained no detectable iSNVs, and the median count was 4 (**S1 Fig**). We found no statistically significant difference in iSNV counts between individuals reporting recent influenza vaccination and those reporting no recent influenza vaccination (**S1B Fig**). Notably, 2 of 283 samples were clear outliers, each with more than 20 iSNVs. One sample (Ct 28.67) from the 2017–18 A(H3N2) season contained 28 iSNVs (3–12.5%) across HA, NA, NP, PA, PB1, and PB2 (**S1 Table**). Notable among these was an iSNV in HA causing an M168I change in antigenic site A at 6.73% (H3 numbering; see Methods). A sample from the 2018–19 A(H3N2) season (Ct 29.66) contained 62 iSNVs (3–5.6%) across HA, PA, PB1, and PB2 (**S1 Table**), of which 54 were present in HA (26 non-antigenic, 28 antigenic). Fifteen of the 28 antigenic iSNVs in HA were nonsynonymous, including antigenic site B mutations S159Y and K160T. Of the 1226 iSNVs detected in our 283 samples, we identified just 64 iSNVs in classical HA antigenic sites (40 nonsynonymous, 24 synonymous) in 31 samples. Taken together, our data support the current consensus that most within-host variants detected during acute influenza virus infections are present at low frequencies, and most samples contain only a few iSNVs—although occasional outliers exhibit substantially greater counts.

**Fig 1.**
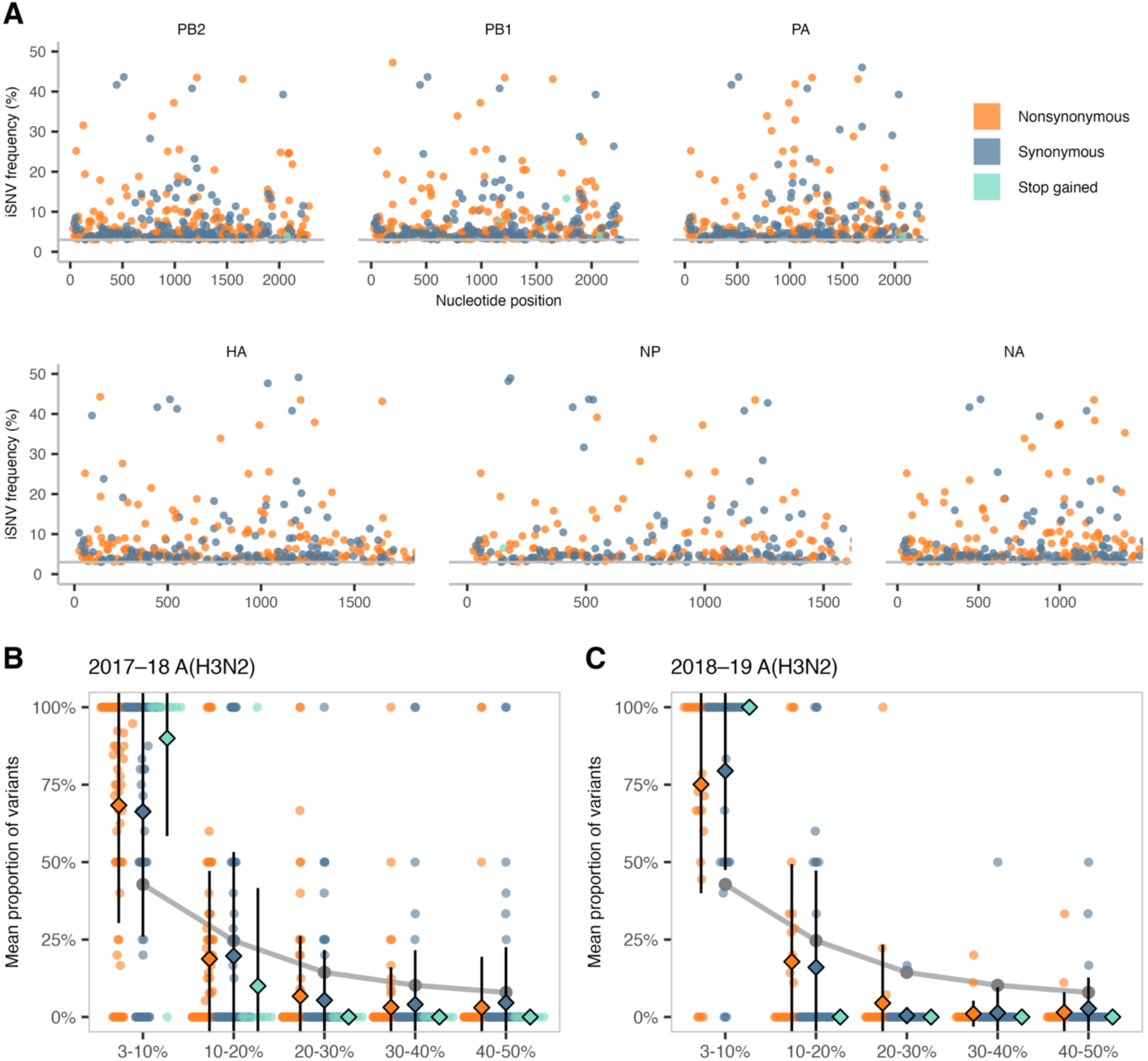
Low-frequency within-host variants predominate across influenza A/H3N2 genes and seasons. **(A)** Intra-host single-nucleotide variants (iSNVs) in at least 3% of sequencing reads (horizontal grey line) in a given sample were plotted for 2017–19 A(H3N2) according to their nucleotide position and frequency. Each dot represents an iSNV colored by mutation type: nonsynonymous (orange), synonymous (blue), or stop gained (green). Genes shown: PB2 (polymerase basic 2), PB1 (polymerase basic 1), PA (polymerase acidic), HA (hemagglutinin), NP (nucleoprotein), and NA (neuraminidase). Not included: MP (matrix; M1/M2) and NS (nonstructural; NS1/NEP) due to overlapping reading frames. **(B** and **C)** iSNV frequency spectra showing the mean proportion of each mutation type for each bin for 2017–18 A(H3N2) and 2018–19 A(H3N2) samples. Error bars denote the standard deviation for each bin’s mutation type. The grey line indicates the proportion of iSNVs in each frequency bin expected under genetic drift. Per-sample proportions are shown as dots for each frequency bin.

We next asked which evolutionary force(s) were most likely to be shaping viral evolution in these acutely infected individuals, as we have done previously [24,25]. Most minority iSNVs are detected below 10% frequency, as shown for A(H3N2) in the 2017–18 (**Fig 1B**) and 2018–19 (**Fig 1C**) seasons. Indeed, iSNVs detected below 10% frequency accounted for a greater proportion of all iSNVs than would be expected under neutral evolution (genetic drift), while, as expected, iSNVs that created stop codons were detected at very low frequencies (15 iSNVs at 3.0–7.5% and one iSNV in PB1 at 13.3%). Such stop-gained iSNVs were detected in PB2, PB1, NP, PA, and NA, but not HA. We next counted the cumulative number of iSNVs for all samples from 3–100% to assess the distribution of iSNVs across various allele frequency bins. Cumulative iSNV counts similarly show that most mutations are present at either very low or very high frequencies within hosts (**S1 Fig**). Together, these results suggest that influenza virus diversity is limited within acutely infected hosts and characterized by weak purifying selection.

### Most iSNVs within hosts are synonymous

To quantify the synonymous and nonsynonymous nucleotide diversity in viral populations within hosts, we calculated *π*, the mean pairwise differences per nucleotide site for a set of sequences [26]. We use *π* here because its value is less sensitive to sequencing read depth than other common diversity metrics [27]. Our analysis revealed that synonymous diversity (*π_S_*) was significantly greater than nonsynonymous diversity (*π_N_*) in every viral gene, both when we considered all seasons and subtypes together (**Fig 2A**; *p* < 0.001, Wilcoxon signed-rank tests) and when we analyzed each influenza season and subtype independently (**S2A–D Fig**; *p* < 0.015). We next sought to quantify the magnitude and directionality of selection occurring within hosts by calculating the difference between *π_N_* and *π_S_* for each gene. The distribution of *π_N_* – *π_S_* values was slightly, but significantly, less than zero in all viral genes, both when we considered all data together across season and subtype (**Fig 2B**; *p* < 0.001, one-sample Wilcoxon signed-rank tests) and also when we analyzed each season and subtype individually (**S2E–H Fig**; *p* < 0.015). Together, these findings suggest that influenza A viruses are generally subjected to purifying selection within acutely infected hosts across influenza seasons, subtypes, and host vaccination status. This is also consistent with the finding that the preponderance of iSNVs are present below 10% frequency (**Fig 1**).

**Fig 2.**
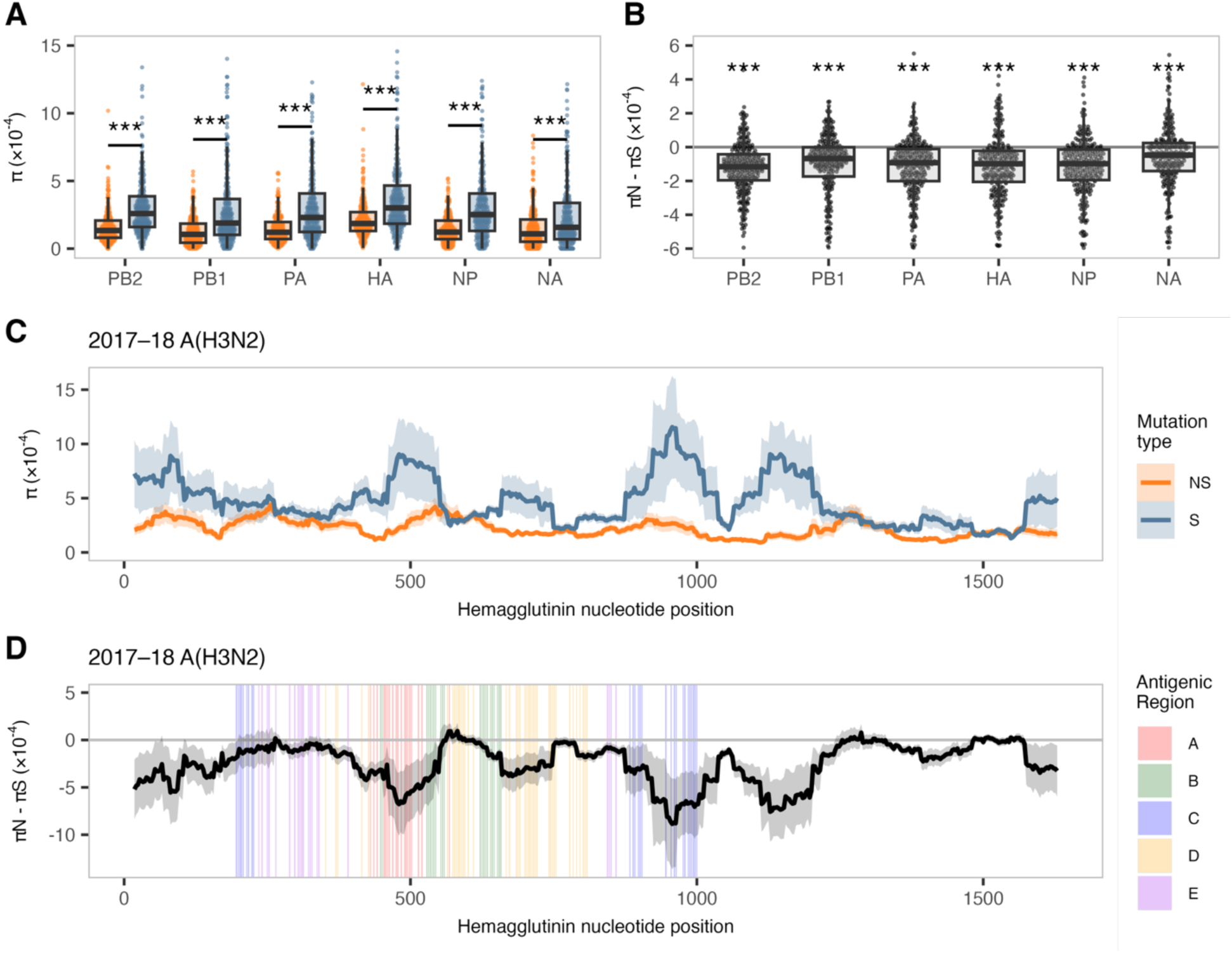
Synonymous diversity exceeds nonsynonymous diversity with evidence of purifying selection. **(A)** The average number of pairwise nonsynonymous differences per nonsynonymous site (*π_N_*, orange) and synonymous differences per synonymous site (*π_S_*, blue) across all analyzed viral genes, subtypes, and seasons. Boxplots are overlaid to indicate the median, interquartile range, and values within 1.5× the interquartile range. **(B)** The difference between *π_N_* and *π_S_* across all viral genes, subtypes, and seasons. Boxplots are overlaid as described in A. The horizontal grey line represents *π_N_* = *π_S_*, with values above zero suggesting diversifying selection and values below zero suggesting purifying selection. **(C)** Mean *π_N_* (orange) and *π_S_* (blue) values for 30-codon sliding windows in the hemagglutinin gene for 2017–18 A(H3N2) viruses. **(D)** Mean *π_N_* – *π_S_* values for 30-codon sliding windows in the hemagglutinin gene for 2017–18 A(H3N2) viruses, colored by antigenic region. Shaded areas denote the standard error of the mean for each codon position. Genes shown: PB2 (polymerase basic 2), PB1 (polymerase basic 1), PA (polymerase acidic), HA (hemagglutinin), NP (nucleoprotein), and NA (neuraminidase). Not included: MP (matrix; M1/M2) and NS (nonstructural; NS1/NEP) due to overlapping reading frames. Paired comparisons of *π_N_* and *π_S_* were tested in A using Wilcoxon signed-rank tests. Median differences between *π_N_* and *π_S_* were tested in B against zero using one-sample Wilcoxon signed-rank tests. Asterisks indicate significance thresholds: *p* < 0.05 (*), *p* < 0.01 (**), and *p* < 0.001 (***); ns = not significant.

Calculating nucleotide diversity for an entire gene could mask selection directed at a small number of specific sites. We focused on HA to address the hypothesis that pre-existing antibody responses may exert selective pressure on viruses that could be detected as elevated *π_N_* – *π_S_* values in certain antigenic regions. We calculated *π_N_* and *π_S_* (**Fig 2C**), as well as the difference between *π_N_* and *π_S_* (**Fig 2D**), using 30-codon sliding windows in HA for each sample. Our analysis of antigenic and non-antigenic regions (defined in H3 as antigenic regions A-E [28,29] and in H1 as sites Ca, Cb, Sa, Sb [30]) revealed that, while values for both *π_N_* and *π_S_* fluctuate along the length of the HA gene, *π_S_* exhibits larger fluctuations. Local spikes in *π_S_* lead to larger (i.e., more negative) *π_N_* – *π_S_* differences in locations that include both classical antigenic sites and non-antigenic regions. This trend is present, with somewhat different patterns, in each influenza season and subtype (**S3 Fig**). Together, these findings show that, although HA genes are under overall purifying selection, the magnitude of purifying selection may vary over the length of HA as well as with season and viral subtype.

### Tight bottlenecks and weak purifying selection shape between-host influenza virus evolution

Our study includes 36 households comprising 46 putative transmission events, which enables us to compare differences in virus population diversity between donors and recipients and test for evidence of selection during transmission. The first individual to report symptoms was identified as the donor case for each household in our dataset. Every other household member who had RT-PCR-confirmed influenza infection within ten days of the donor case was identified as a potential recipient case. Prior work has noted that household members might coincidentally acquire independent infections from outside the household [31–33], potentially confounding analyses of transmission events. To address this, we implemented a genetic criterion for household pairing, as described previously [24]. Briefly, we calculated the proportion of all iSNVs shared between each donor-recipient pair and excluded pairs with shared proportions below the 95th percentile of those observed in random community pairs (**Fig 3A**). The random community pairing followed the expected distribution: 95% of random community pairs shared fewer than 11.2% of variants (vertical dashed line). That is, 40 of the 46 potential household pairs shared a higher proportion of variants than 95% of randomly paired community samples. As a complementary approach to determine the relatedness of viruses infecting members of the same household, we generated maximum likelihood phylogenies for the HA consensus sequences of each influenza season and subtype (**S4 Fig**). Viruses from members of the same household clustered closely together on all trees, with identical or near-identical consensus sequences. Together, our epidemiological and genetic criteria enable us to rigorously identify cases of within-household transmission.

**Fig 3.**
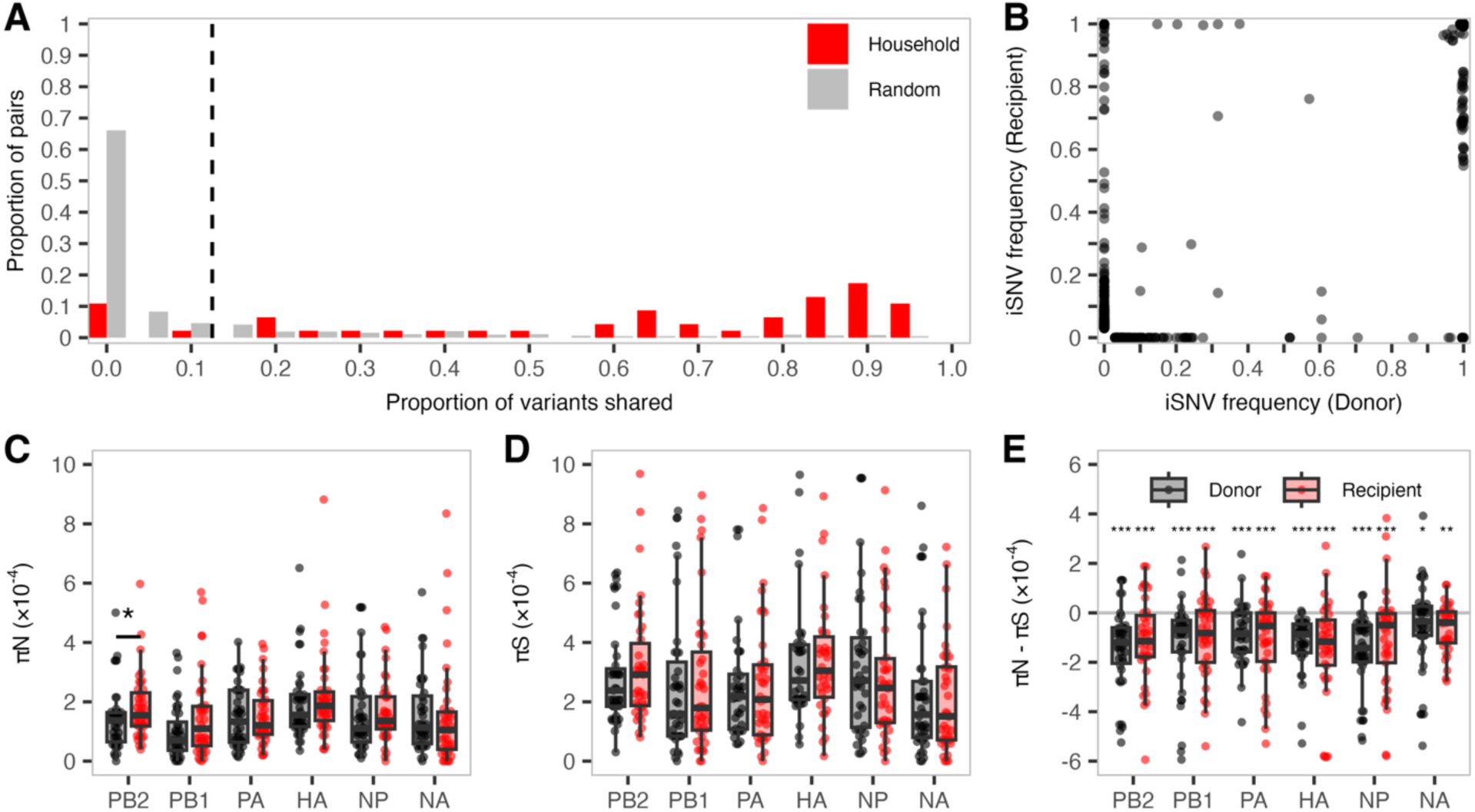
Donor–recipient variant frequencies and nucleotide diversity across genes. **(A)** The proportion of variants shared between putative household transmission pairs (red) or randomly paired virus sequences from a given season and subtype (grey) for all samples included in this study. The vertical line denotes the threshold below which 95% of random community pairs fell. **(B)** iSNV frequency in all donors and recipients as defined in A. Frequencies are shown as 0% in the paired individual if the iSNV was absent or below the 3% quality-control threshold. Boxplots are overlaid to indicate the median, interquartile range, and values within 1.5× the interquartile range for **(C)** *π_N_*, **(D)** *π_S,_* and **(E)** *π_N_* – *π_S_* values for all donor (black) and recipient (red) samples across all viral genes analyzed. Genes shown: PB2 (polymerase basic 2), PB1 (polymerase basic 1), PA (polymerase acidic), HA (hemagglutinin), NP (nucleoprotein), and NA (neuraminidase). Not included: MP (matrix; M1/M2) and NS (nonstructural; NS1/NEP) due to overlapping reading frames. The horizontal grey line in E represents zero, where *π_N_* = *π_S_*. Paired comparisons of nucleotide diversity between donors and recipients were tested in C and D using Wilcoxon signed-rank tests. Median differences between *π_N_* and *π_S_* were tested in E against zero using one-sample Wilcoxon signed-rank tests. Asterisks indicate significance thresholds: *p* < 0.05 (*), *p* < 0.01 (**), and *p* < 0.001 (***); ns = not significant.

We next determined how transmission affected iSNV frequencies in the donor-recipient pairs. If transmission bottlenecks are loose, we expect iSNV frequencies in the recipient to resemble those in the donor. In total, we detected 1330 iSNVs in either donors or recipients. We categorized these iSNVs into three groups: present in both donor and recipient, present only in the donor, or present only in the recipient. The majority (72%) of iSNVs were detected in both the donor and recipient; of these, 99% exceeded 50% frequency (consensus level) and 95% were present above 90% frequency in both the donor and recipient. In contrast, only eight iSNVs (five synonymous, three nonsynonymous) below 90% were observed in both the donors and recipients at similar frequencies. These results indicate that highly abundant donor iSNVs compose most of the diversity shared between donors and recipients.

In one household pair, a synonymous substitution in PB1 codon 346 (PB1: N346N) was observed at 57% frequency in the donor and 76% in the recipient (**Fig 3B**, **S2 Table**). In a separate household pair, we detected a synonymous PB1 substitution (PB1: P627P) at 10% frequency in the donor and 29% in the recipient (**S2 Table**). In a third household cluster, a nonsynonymous PB2 substitution (PB2: A270S) was present at 60% frequency in the donor and 0%, 6%, and 15% in the three recipients (**S2 Table**). In a fourth household cluster, three intermediate-frequency iSNVs were potentially transmitted from the donor to three recipients (**S2 Table**). The donor sample harbored two NP iSNVs: T22A at 24% and G187G at 10%. Both mutations were lost in Recipient 1, while only T22A persisted at 30% in Recipient 2, and only G187G was maintained at 15% in Recipient 3. Similarly, in the same household, a synonymous HA mutation (HA: C471C, H1 numbering) was found at 32% in the donor, but was present at 100%, 71%, and 14% in the three recipients. Conversely, a nearby synonymous HA mutation (HA: G463G, H3 numbering) was present at 14% in the donor but was undetected in all recipients. The fact that intermediate-frequency donor iSNVs are lost, maintained, and/or fixed in different patterns in different recipients of the same donor is consistent with the idea that transmission of seasonal influenza A viruses from acutely infected humans typically involves a very tight, stochastic bottleneck in agreement with previous reports [7,20].

Twelve percent of iSNVs were only present in the donor, with 91% of these detected below 50% frequency. Meanwhile, 16% of iSNVs were detected only in the recipient, and 89% of these recipient-only iSNVs were minority variants. We cannot determine whether such iSNVs were present in the donor below our detection limits or whether they emerged *de novo* in the recipient. Notably, 7% of the recipient-only iSNVs (representing 1% of all iSNVs in donors and recipients) were fixed in the recipient despite being undetected in the donor. Of these fixed recipient-only iSNVs, 4 were synonymous and 7 were nonsynonymous. Together, our results suggest that transmission of seasonal influenza A viruses generally involves very tight bottlenecks in which the likelihood that an iSNV is transmitted depends largely on its frequency in the donor.

While substantial within-host diversity is lost during transmission, the extent to which selection shapes this reduction remains unclear. We therefore next compared *π_N_* and *π_S_* between donors and recipients to assess whether transmission bottlenecks preferentially restrict nonsynonymous mutations. Overall, *π_N_* and *π_S_* values did not change significantly during transmission in any viral genes (**Fig 3CD**), except in PB2, which showed slightly but significantly higher *π_N_* values in recipients relative to donors (*p* = 0.025). This significant increase was driven by higher *π_N_* values specifically in the 2017–18 A(H3N2) group (*p* = 0.003; **S5A Fig**). In the 2017–18 A(H3N2) group, we also observed several additional genes in which recipients had slightly but significantly higher *π_N_* values compared to donors (PA and HA: *p* < 0.05), along with a modest increase in *π_S_* values for PB2 and NA (*p* < 0.05). These patterns were not observed in other groups (**S5 Fig**, **S6 Fig**). Across all genes when aggregating all seasons and subtypes in this study, the distribution of *π_N_* – *π_S_* values was significantly below zero (*p* < 0.05), further suggesting that purifying selection was the dominant selective force in donors and recipients (**Fig 3E**). Statistical comparisons between donors and recipients in **Fig 3E** yielded no significant differences (**S3 Table**). Taken together, our results indicate that, on balance, purifying selection appears to be equally strong in both donors and recipients.

### Antigenic sites on hemagglutinin are under distinct selective pressures during transmission

The model of influenza virus antigenic evolution proposed by Morris *et al.* [19] posits that selection operates mainly at the time when a new infection is established. Because this is expected to be a rare event, detecting it would require access to many well-defined transmission pairs. To ascertain whether this phenomenon was detected in our donor-recipient pairs (**Fig 3B**), we investigated whether specific nonsynonymous mutations in antigenic regions of HA were acquired or increased in frequency during transmission. We detected the transmission of a nonsynonymous antigenic site variant in just one transmission pair, in which both the donor and recipient were vaccinated. A nonsynonymous mutation encoding HA I185T (H1 numbering, antigenic site Sb of 2018–19 A(H1N1)pdm09) became fixed across transmission (from 38% in the donor to 100% in the recipient; **S2 Table**). This event is consistent with selection during transmission; however, it could also have become fixed because of the tight transmission bottleneck and not because it conferred a fitness benefit.

Four other transmission pairs maintained HA I185T at 100% frequency, and this mutation was present in 19 other samples from our cohort at 100% frequency, suggesting that it was circulating in the community during the study period. The four donor samples contained an additional 23 mutations in HA, which were fixed relative to the within-season reference, including 2 synonymous mutations in antigenic region Ca1 (HA: E235E and HA: G237G). All of these mutations were present in the recipients, indicating that viruses infecting these donor-recipient pairs were divergent relative to most viruses sampled from the community during this time period (**S4D Fig**, taxon boxed in red). In all other donor-recipient pairs, we saw no stark increase in the frequency of iSNVs in classical antigenic sites, consistent with the expectation that this phenomenon is infrequent.

Although there was only one instance of an iSNV with potential antigenic impact increasing in frequency during transmission, we reasoned that patterns of nucleotide diversity could indicate whether selection acted specifically on antigenic sites of HA. To test this, we quantified the difference between *π_N_* and *π_S_* values for each codon in antigenic and non-antigenic regions (defined in H3 as antigenic regions A-E [28,29] and in H1 as sites Ca, Cb, Sa, Sb [30,34]) in each donor and recipient sample, then we normalized per-sample differences by the number of sites in each region. Across donors, recipients, and the pooled “all” group (i.e., all samples in the dataset considered together), antigenic and non-antigenic regions displayed significantly negative per-site *π_N_* – *π_S_* values (*p* < 0.0231), consistent with purifying selection (**Fig 4**). These values remained stable during transmission, with no significant differences between donors and recipients for either region—a pattern that held across all seasons and subtypes analyzed in this study (**S7 Fig**). We detected a significant difference between antigenic and non-antigenic regions only in the “all” group (*p* = 3.69e-11), where non-antigenic regions showed stronger purifying selection. In this group, the median per-site *π_N_* – *π_S_* value for antigenic regions was zero, with 32% of values exactly zero and 40% falling below zero. These results indicate that purifying selection is generally weaker in antigenic regions of HA than in non-antigenic regions. Still, it remains unclear whether this reflects reduced constraint, increased selective pressure, or both. If distinct selection pressures do act on antigenic sites during transmission, they appear too weak or rare to detect with this approach.

**Fig 4.**
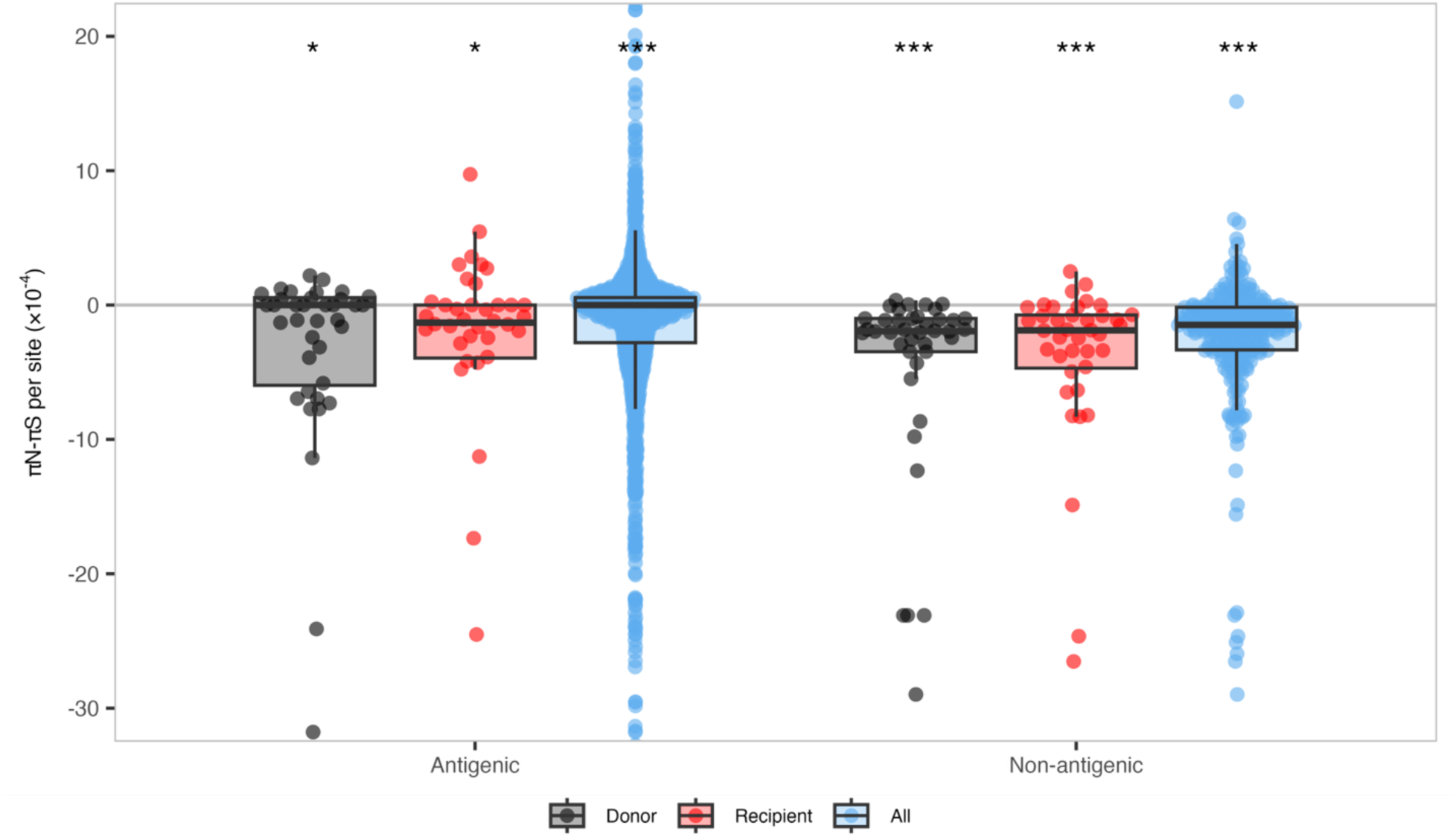
Nucleotide diversity across antigenic and non-antigenic regions of HA. Per-site *π_N_* – *π_S_* values in antigenic and non-antigenic regions of HA for all samples in this study. *π_N_* – *π_S_* values were normalized to account for differences in the amount of antigenic and non-antigenic sites, denoted *π_N_* – *π_S_* per site. Boxplots are overlaid to indicate the median, interquartile range, and values within 1.5× the interquartile range for each group: Donor (black), Recipient (red), or All (blue). The “All” group includes values for all viral genomes sampled (donors, recipients, and also those without a linked household contact). The horizontal grey line represents zero, where *π_N_* = *π_S_*. Median differences between *π_N_* and *π_S_* were tested against the null hypothesis (*π_N_* – *π_S_* = zero) using one-sample Wilcoxon signed-rank tests. Asterisks indicate significance thresholds: *p* < 0.05 (*), *p* < 0.01 (**), and *p* < 0.001 (***); ns = not significant.

## Discussion

On the scale of the human population, influenza viruses clearly evolve under immune selection pressure. However, we are confronted by a paradox of scale: new antigenic variants must arise within individual infected hosts, but viral evolution within and between infected hosts appears to be dominated by randomness and purifying selection, making it difficult for new variants to emerge and transmit. Defining the evolutionary mechanisms connecting the individual to the global scale will enhance our ability to understand and forecast trends in influenza virus antigenic evolution [8]. Morris *et al.* predict that selection occurs not over time within hosts, but during the establishment of a new infection [19]. Here, we used 283 high-quality influenza A-positive samples from a community cohort study. Of these, 71 samples formed 40 validated donor–recipient pairs across 31 households for transmission analyses, and the remaining 212 single infections informed within-host analyses. We used these pairs to assess whether there was evidence for selection during transmission. Our study supports previous findings suggesting that the short duration of acute influenza virus infection, coupled with purifying selection and anatomical compartmentalization, dampens the generation of substantial diversity within hosts [6–8,17,27,35–37]. We also find that narrow transmission bottlenecks between hosts result in a substantial loss in low-frequency diversity and dampen the ability of selection to act efficiently during transmission.

Mutations arise during acute influenza virus infection and are culled or amplified by purifying or diversifying selective pressures, respectively. Consistent with previous studies, we observe low-frequency iSNVs across the viral genome (**Fig 1**) and signals of purifying selection on every influenza virus gene (**Fig 2B**) [7]. To become dominant in human populations, variants must survive transmission bottlenecks and propagate in additional hosts. If transmission bottlenecks were extremely loose (allowing many viruses to be transferred from donor to recipient) and under neutral selection, we would expect iSNVs to be present in the donor and recipient at similar frequencies. Conversely, under a tight bottleneck, low-frequency iSNVs are rarely transmitted, but if they are, the limited number of unique founding genomes often causes these variants to be present at high frequency in the recipient. In the 31 households and 40 transmission events we analyzed, low-frequency donor iSNVs were generally lost, and the vast majority of transmitted iSNVs were ≥90% frequency in the recipient. This pattern is consistent with a narrow, stochastic bottleneck, as others have reported previously for acute transmission of influenza A and SARS-CoV-2 [7,24,25,37,38].

Importantly, selection for specific variants during transmission may also result in reduced viral genetic diversity in recipients in a manner that may be difficult to distinguish from a stochastic bottleneck [39]. We have observed such selective bottlenecks ourselves in experimental systems of influenza A virus transmission in ferrets [4,21]. However, in the present work, our data are mostly consistent with the stochastic model: we see dramatic shifts in iSNV frequency, in which iSNVs are more likely to transmit if they are at high frequency in the donor. There is no single iSNV that is consistently transmitted. In several households, we capture transmission from a single donor to multiple recipients, and in these cases, we see that each recipient acquired a different combination of iSNVs. This suggests that the iSNVs are present in different viruses (i.e., not genetically or physically linked) and that each recipient received a random sample of viruses present in their donor.

We expect selective bottlenecks to be rare in acute human influenza transmission, but it is interesting that we did observe one nonsynonymous mutation in an HA antigenic region, HA: I185T, which is in antigenic site Sb. This mutation was a minority variant in the donor and became fixed in the recipient. Fixation of a potential antigenic variant during transmission could be consistent with the Morris *et al.* hypothesis [19], but it is also important to consider the broader context of contemporaneous viral diversity: A(H1N1)pdm09 viruses circulating both globally and in the sampled community were polymorphic at HA:185 site during the study period. Threonine (T185) had been dominant at this position, but isoleucine (I185) was becoming more frequent in the 2018–19 season, present in 51% of our samples and 66% of global sequences (**S8 Fig**) [40,41]. The donor in our transmission pair carried 185T at 38% frequency and 185I at 62%, yet the recipient became productively infected with 185T alone, the minority variant of the donor and the globe at the time. Thus, given the cocirculation of both T185 and I185 in our community and our lack of information about immune responses to influenza A viruses in our cohort, we cannot determine whether transmission of this mutation was favored by natural selection in this donor-recipient pair.

Our findings are consistent with previous studies suggesting that tight transmission bottlenecks reduce the ability of selection to fix beneficial iSNVs during transmission [7,20,21,37,38]. When selection is weak or absent, the probability that an iSNV is transmitted will be a function of its frequency rather than fitness. To test whether selection might still act preferentially on certain regions, we compared levels of purifying selection across antigenic and non-antigenic sites of HA. As a whole, both antigenic and non-antigenic regions of HA were under purifying selection, and our analysis detected no significant difference in purifying selection on antigenic or non-antigenic regions between donors and recipients (**Fig 4**). Non-antigenic regions showed slightly stronger purifying selection than antigenic regions, but this difference was not significant across most comparisons. Overall, within hosts, antigenic regions of HA appear to be under very weak purifying selection, while non-antigenic regions experience slightly stronger, yet still weak, purifying selection. This pattern was not clearly observed during transmission. While we did not observe any striking differences between the antigenic evolution of A(H1N1)pdm09 and A(H3N2), we are limited by the relatively small number of annotated antigenic sites for these viruses (50 and 130, respectively). Our results further support the idea that stochastic processes dominate influenza A virus evolutionary dynamics during transmission and that selective transmission leading to the global emergence of HA antigenic variants is an extremely rare, high-consequence event that will be difficult to observe in nature.

Our study has several limitations. We were unable to perform deep sequencing in technical replicates for this work, which could reduce the accuracy of iSNV frequencies, particularly near the 3% frequency threshold. To account for this deficit, we implemented stringent quality control criteria to eliminate the inclusion of samples of insufficient quality to detect low-frequency variants. Further, we only included samples with high whole-genome coverage and depth to facilitate comparison between genes. Our findings are also partially limited in that we have excluded M1/M2 and NS1/NEP genes from diversity analyses due to their overlapping reading frames and small size, which can confound site-based diversity metrics [8]. Our study design has additional limitations in both demographic and sampling structure. By nature of the study design, school-aged children reporting respiratory symptoms are the first household members enrolled in each household, and as a result, are likely to be recognized as donors in our transmission study. Indeed, all donors in our cohort’s transmission networks were children. This raises a potential bias, as children may have had more recent or limited exposure to influenza viruses compared to adult household members, potentially affecting their within-host diversity [31,42,43]. The study design also means that our cohort may be biased toward symptomatic infections in the donors. Importantly, we must caution that we do not know precisely when transmission occurred between putative donors and recipients. As such, donor samples may be derived later in the donor’s acute infection than recipient samples. Additionally, misclassification of donor-recipient direction remains possible, particularly when household members have shared community exposure or separate introductions of genetically similar viruses. The key limitation of our study is our relatively few validated transmission events. For future studies to rigorously evaluate the possibility that selection for antigenic variants occurs during transmission, larger cohorts will be required that focus on enrolling households or people in other settings where transmissions are likely to be captured. Future studies should employ similar prospective cohort study designs and analyze deep-sequencing efforts alongside immunological and epidemiological data to expand upon our findings.

## Materials and Methods

### ORCHARDS participant recruitment and sampling

The full Oregon Child Absenteeism Due to Respiratory Disease Study (ORCHARDS; Oregon, Wisconsin, USA) methodology can be found in Temte *et al.* 2022 [23]. Briefly, ORCHARDS was a community cohort study designed to evaluate usability of cause-specific student absenteeism as a method for early detection of increased influenza activity in schools and surrounding community; ORCHARDS also served as a platform to study the characteristics and impact of different respiratory pathogens affecting school-aged children. A subset of participants in this cohort was recruited into a sub-study to assess within-household transmission of respiratory pathogens. Household members were responsible for collecting specimens on Day 0 (within 24 hours of the home visit by study coordinators) and Day 7 (seven days after the initial collection). Each household participant collected the specimen without staff observation on the day of the home visit (Day 0) and again seven days later (Day 7). Two participants deviated from this protocol: one collected their first sample on Day 1 and another collected their second sample on Day 8. Both samples were retained for analysis (**S9 Fig**). In total, this study provided us with access to specimens from 384 children that tested positive for influenza A virus by RT-PCR (IVD CDC Human Influenza Virus RT-PCR Diagnostic Panel [44]) using the Applied Biosystems™ 7500 Real-Time PCR instrument (Thermo Fisher) at the Wisconsin State Laboratory of Hygiene (WSLH).

### Illumina library preparation and sequencing

Following the manufacturer’s protocol, influenza virus RNA was extracted from nasal swabs using the QIAcube HT (Qiagen) with the QIAamp 96 Virus QIAcube HT Kit (Qiagen). RT-PCR was then performed on extracted viral RNA using the SuperScript™ III One-Step RT-PCR System with Platinum™ Taq High Fidelity DNA Polymerase (Thermo Fisher) using the primers described by Zhou *et al.* [45], which amplify all of the eight influenza A virus gene segments in a single RT-PCR reaction, with the following thermal cycling conditions: 42°C for 50 min, 50°C for 10 min, 94°C for 2 min, four cycles of the following: [94°C for 30 sec, 43°C for 30 sec, 68°C for 3 min 50 sec], 30 cycles of the following: [94°C for 30 sec, 57°C for 30 sec, 68°C for 3 min 30 sec increasing by 10 sec every cycle], then finally 68°C for 10 min. The resulting PCR amplicons are then treated with Exonuclease I (Thermo Fisher) and incubated at 37°C for 15 min and 80°C for 15 min. The DNA is then quantified using Victor X2 (Perkin Elmer) using the Quant-iT High-Sensitivity dsDNA kit (Thermo Fisher) and normalized to 0.2 ng/μL. The DNA library is prepared for sequencing using the Nextera XT DNA Library Prep Kit (Illumina) following the manufacturer’s protocol and sequenced using the MiSeq platform (Illumina).

### iSNV calling pipeline

Intra-host single-nucleotide variants (iSNVs) were called using bespoke scripts through the Center for High Throughput Computing (CHTC) at the University of Wisconsin–Madison. All read processing scripts were executed through the rieshunter/tcflab:v2.04 Docker image (https://hub.docker.com/r/rieshunter/tcflab/tags) on a CHTC server node. This Docker image consists of a Linux GNU OS (18.04.6 LTS “Bionic Beaver”) with all necessary commands and program files (Docker version 20.10.13).

Raw, paired FASTQ files from samples positive for the subtype of interest were adapter-trimmed and quality-trimmed to an average 5-base sliding-window quality of Q20 using Trimmomatic (version 0.39). Additionally, all reads less than 100 bases in length were discarded. Trimmed, paired reads were then merged using BBMerge.sh (BBMap version 38.96), outputting merged pairs and unmerged files. Merged and unmerged reads were normalized to 2000 reads for all 31-mers above 200 depth. Normalized, merged, and unmerged reads were aligned to the season-appropriate vaccine influenza strain, determined by sample collection time and PCR subtyping, using bwa mem (BWA version 0.7.17). Each sample was aligned to its respective season- and subtype-specific reference vaccine strain: A/Hong Kong/4801/2014 (2017–18 A(H3N2)), A/Singapore/INFIMH-16-0019/2016 (2018–19 A(H3N2)), or A/Michigan/45/2015 (2017–19 A(H1N1)pdm09) [44,46]. Alignments were concatenated with samtools merge to produce a single sample alignment file from the merged file and the R1 and R2 unmerged files (Samtools version 1.15). iSNVs were called using callvariants.sh (BBMap version 38.96) to a minimum Phred quality score of Q30 and minimum position coverage of 100. A minimum frequency of 3% was applied after post-processing analysis in RStudio.

To generate consensus sequences for each sample, variants above 50% were applied to each sample’s respective season- and subtype-specific reference vaccine strain sequence using BCFtools (version 1.15). Samples that failed to map to one or more gene segments, had <100x coverage at one or more segments, or <99% genome coverage were discarded. High-quality sample consensus genomes were split by segment and aligned to one another using ClustalO (version 1.2.4), and these alignments were visualized with JALview (version 2.11.2.6) to generate a single within-season consensus sequence for each season-subtype combination.

We then realigned all normalized reads to their respective within-season reference sequence to contextualize iSNVs relative to circulating strains in the community. Alignment and variant-calling were performed using the same quality metrics above to produce “realigned” alignment files and “recalled” iSNV files. Samples failing whole-genome coverage thresholds (described above) after realignment were excluded from downstream analyses. To detect and remove potential outliers with excess within-host diversity, we quantified the number of iSNVs >0.5% allele frequency for each sample, ranked them in descending order within their season-subtype group, and excluded samples that were in the top 10%, as in Xue *et al.* [8].

### Diversity statistics

Variants were annotated using snpEff (version 5.1) relative to the within-season reference sequence to categorize mutation type (synonymous, nonsynonymous, stop-gained, etc.). Antigenic regions were defined in H3 as antigenic regions A-E [28,29] and in H1 as sites Ca, Cb, Sa, Sb [30,34]. Amino acids composing antigenic sites H3 and H1 HAs are listed in **Supplemental Tables 4 and 5**, respectively. The per-segment, per-gene, and per-codon π diversity were calculated with SNPGenie (version 2019.10.31). Sample π diversity was computed as a per-nucleotide weighted average of each segment’s π diversity.

Diversity statistics, data processing, and figure generation were performed with GNU bash (version 3.2.57(1)-release) and R (version 4.4.1 “Race for Your Life”) through macOS Sequoia 15.5. Multiple packages were used in RStudio (version 2023.12.1+402 “Eye Holes”): tidyverse 2.0.0 (which loads tibble 3.2.1, dplyr 1.1.4, purrr 1.0.2, readr 2.1.5, tidyr 1.3.1, stringr 1.5.1, forcats 1.0.0), vcfR 1.15.0 for parsing VCF files, reshape2 1.4.4 for reshaping data, lubridate 1.9.3 for date-time manipulation, ggplot2 3.5.1 for graphics, plus cowplot 1.1.3, gridExtra 2.3, ggpubr 0.6.0, scales 1.3.0, and ggbeeswarm 0.7.2 for figure layout and annotation. All scripts are publicly available via GitHub (https://github.com/RiesHunter/ORCHARDS).

### Statistical comparisons

Statistical comparisons were performed in R using the base-R wilcox.test function. We first assessed normality with the Shapiro–Wilk test: (shapiro.test). For non-normally distributed data, we employed non-parametric statistical tests. Paired Wilcoxon signed-rank tests (wilcox.test(x, y, paired = TRUE)) were used for matched (dependent) samples. The null hypothesis is that the median of the paired differences equals zero. One-sample Wilcoxon signed-rank tests (wilcox.test(x, mu = 0)) were used to test whether the sample median differs from zero. Unpaired two-sample Wilcoxon rank-sum tests (wilcox.test(x, y, paired = FALSE)), also known as Mann–Whitney U tests, were used for unmatched (independent) samples. The null hypothesis is that the two groups have the same distribution. All tests were two-sided, and *p* < 0.05 was considered significant. Statistical test *p*-values for each figure can be found in **Supplemental Table 3**.

### Donor-recipient pairing

Every ORCHARDS sample that passed quality control was paired based on household status and timing between infections. Individuals must have been within the same household to be considered pairs. Further, “donors” must have tested positive on the same day or before the “recipient’s” sample was collected. To improve confidence in household pairing status, all samples within the same season and influenza subtype were randomly paired, and a proportion of shared variants was calculated for each random pair, as we have done previously [24]. This proportion was also calculated for potential household pairs. The proportion of shared variants was calculated as twice the number of shared variants between the two samples divided by the total number of variants between the two samples. Household pairs sharing a higher proportion of variants than 95% of random pairings within the season were considered valid donor-recipient pairs. Validated transmission pairs are shown in **Supplemental Figure 9**.

### Maximum likelihood tree generation

All hemagglutinin consensus sequences were stratified by season and subtype and aligned with ClustalW (version 2.1) [47]. Aligned sequences were the input for RAxML-NG (version 1.2.2) [48] using the GTR+Gamma substitution model. RAxML-NG bestTree files were analyzed in RStudio and R using APE 5.8 [49], phangorn 2.12.1 [50], cowplot 1.1.3, RColorBrewer 1.1-3, ggplot 3.5.1 [51], and ggtree 3.12.0 [52–54]. Briefly, trees were imported with read.tree, roots were resolved against the respective vaccine strain reference sequence, and tips were colored by household.

### Data availability

The consensus genome sequences are available from GISAID (www.gisaid.org). Data, analysis pipelines, and figures are available for replication of these results on GitHub (https://github.com/RiesHunter/ORCHARDS).

## Acknowledgments

We would like to thank the participants from this study as well as the Oregon School District staff, ORCHARDS team, and Wisconsin State Laboratory of Hygiene colleagues for their contribution to this study.

## Disclaimers

The findings and conclusions in this report are those of the authors and do not necessarily represent the official position of the US Centers for Disease Control and Prevention (CDC).

## Disclaimer

The findings and conclusions in this study are those of the authors and do not necessarily represent the official position of the U.S. Centers for Disease Control and Prevention.

## Author contributions

H.J.R. contributed to data curation, methodology, formal analysis, project administration, bioinformatics software, data visualization, writing—draft preparation, and draft review and editing. J.L. contributed to data curation, methodology, bioinformatic software, and writing—draft preparation. K.F. contributed to data curation and methodology.

S.B. contributed to data curation and methodology. M.G. contributed to data curation and methodology. R.G. contributed to data curation and methodology. T.D. contributed to data curation and methodology. K.R. contributed to data curation and methodology. A.U. contributed to conceptualization, data curation, methodology, patient recruitment, and project administration. J.T. contributed to conceptualization, data curation, methodology, patient recruitment, and project administration. T.C.F. contributed to conceptualization, data curation, methodology, project administration, writing—draft review and editing, and supervision.

## Additional information

Correspondence and requests for materials should be emailed to T.C.F.

## Supporting Information

**S1 Fig.**
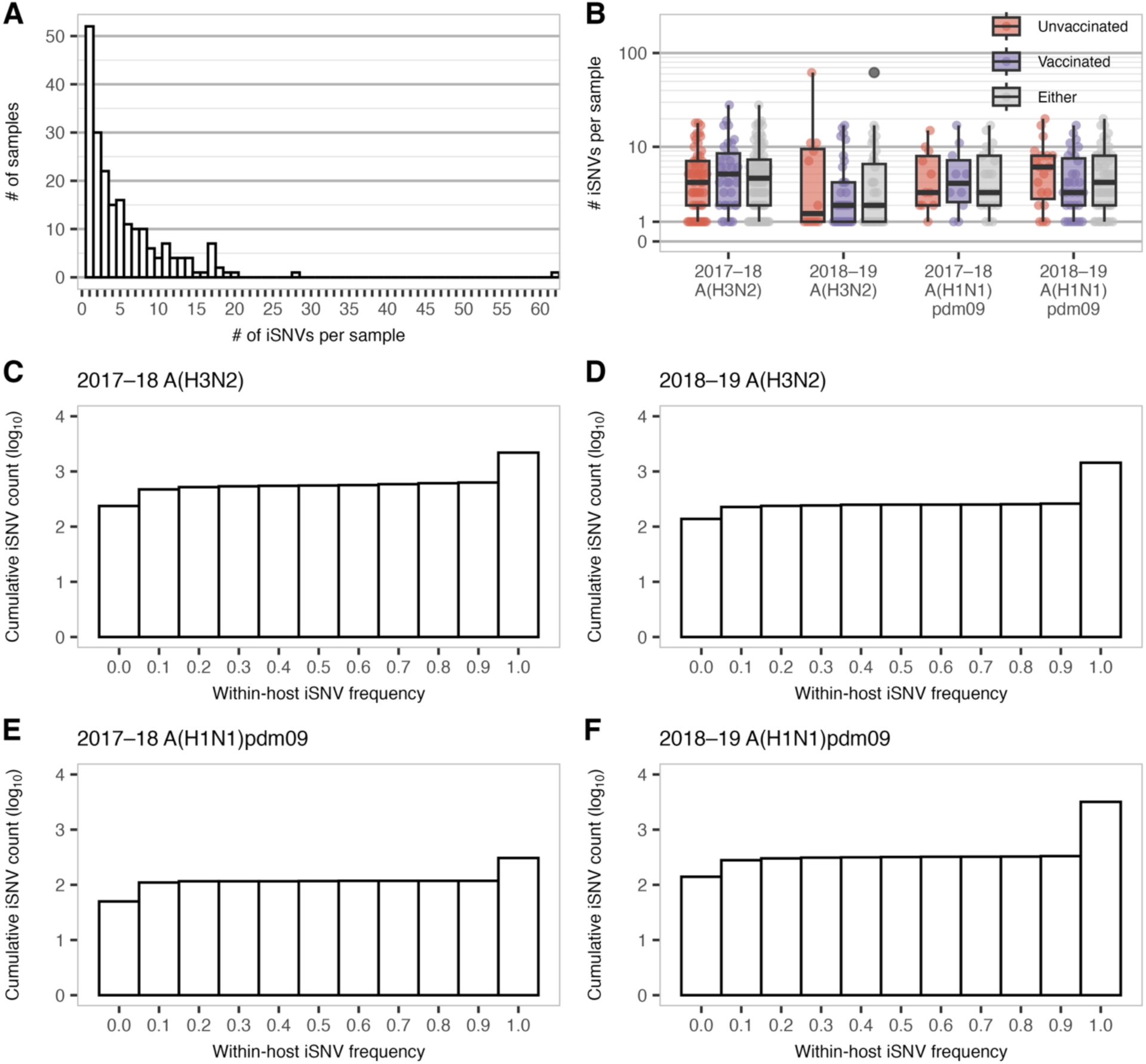
Within-host iSNV counts and cumulative frequency by influenza season and subtype. **(A)** Histogram depicting the number of samples containing the indicated numbers of intra-host single-nucleotide variants (iSNVs) across all genes included in this study. **(B)** Number of iSNVs detected per sample (dots), stratified by reported recent influenza vaccination status: unvaccinated (red), vaccinated (purple). Vaccination refers to current-season influenza vaccination at specimen collection, confirmed in the Wisconsin Immunization Registry [23]. Either (grey) includes all samples regardless of vaccination status. Boxplots are overlaid to indicate the median, interquartile range, and values within 1.5× the interquartile range for iSNV counts. Histograms of the cumulative number of iSNVs by frequency for **(C)** 2017–18 A(H3N2), **(D)** 2018–19 A(H3N2), **(E)** 2017–18 A(H1N1)pdm09, and **(F)** 2018–19 A(H1N1)pdm09. Each bin spans a 10% frequency interval of the form [a, b), where the lower bound is included and the upper bound is excluded—except for the final bin, which includes values equal to 100%. Variants below 3% frequency are excluded due to quality filtering. Paired comparisons of iSNV counts between vaccinated and unvaccinated groups within each season-subtype were tested in B using Wilcoxon rank-sum tests (AKA Mann-Whitney U tests). Asterisks indicate significance thresholds: *p* < 0.05 (*), *p* < 0.01 (**), and *p* < 0.001 (***); ns = not significant.

**S2 Fig.**
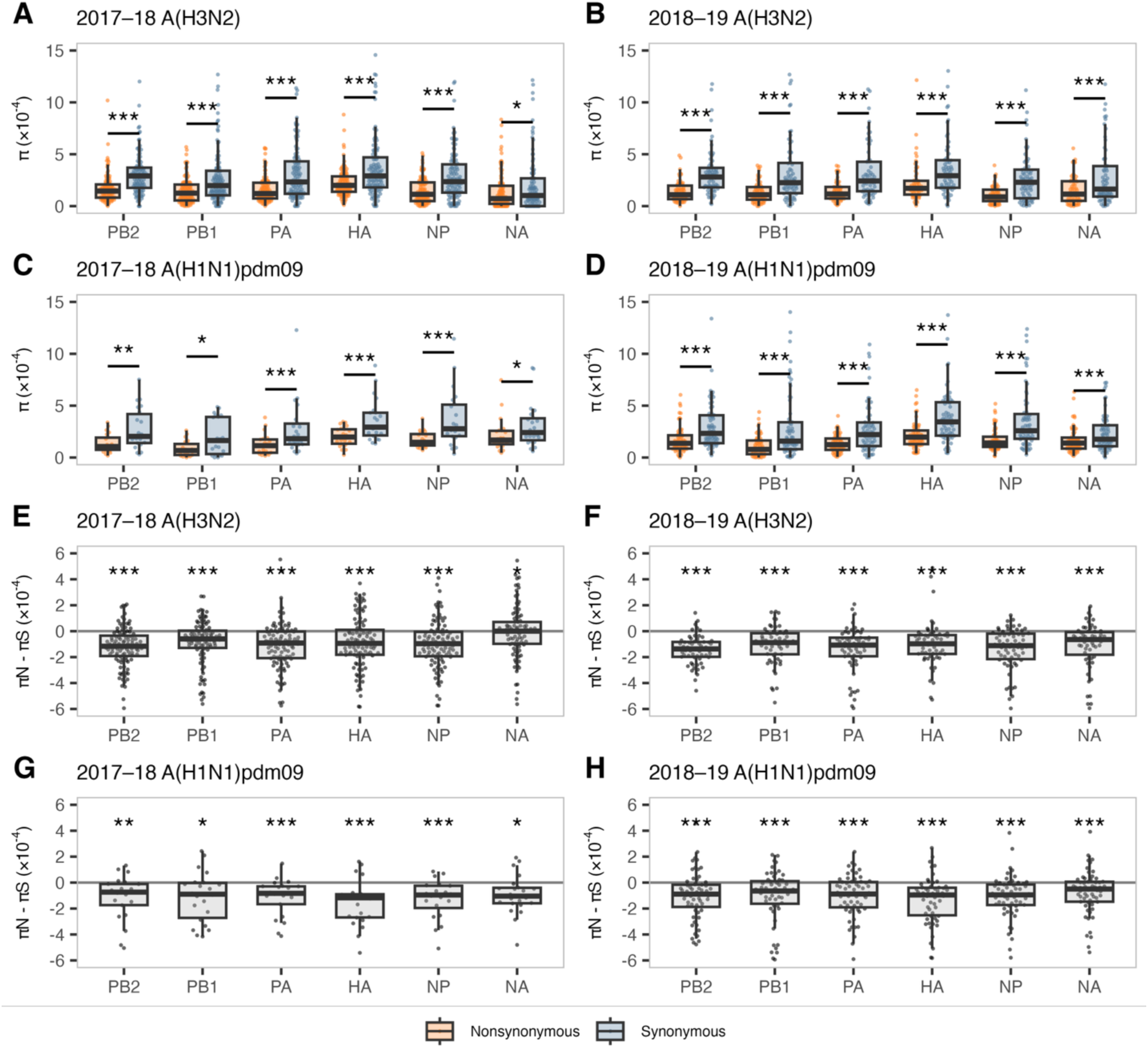
Gene-wise nucleotide diversity across influenza seasons and subtypes. (A–D) The average number of pairwise nonsynonymous differences per nonsynonymous site (*π_N_*, orange) and the average number of pairwise synonymous differences per synonymous site (*π_S_*, blue) across all genes included in this study for **(A)** 2017–18 A(H3N2), **(B)** 2018–19 A(H3N2), **(C)** 2017–18 A(H1N1)pdm09, and **(D)** 2018–19 A(H1N1)pdm09. Boxplots are overlaid to indicate the median, interquartile range, and values within 1.5× the interquartile range. **(E–H)** The difference between *π_N_* and *π_S_* for **(E)** 2017–18 A(H3N2), **(F)** 2018–19 A(H3N2), **(G)** 2017–18 A(H1N1)pdm09, and **(H)** 2018–19 A(H1N1)pdm09. Genes shown: PB2 (polymerase basic 2), PB1 (polymerase basic 1), PA (polymerase acidic), HA (hemagglutinin), NP (nucleoprotein), and NA (neuraminidase). Not included: MP (matrix; M1/M2) and NS (nonstructural; NS1/NEP) due to overlapping reading frames. Boxplots are overlaid as described in above. The horizontal grey line represents *π_N_* = *π_S_*. Paired comparisons of *π_N_* and *π_S_* were tested using Wilcoxon signed-rank tests. Median differences between *π_N_* and *π_S_* were tested against the null hypothesis (i.e., *π_N_* – *π_S_* = zero) using one-sample Wilcoxon signed-rank tests. Asterisks indicate significance thresholds: *p* < 0.05 (*), *p* < 0.01 (**), and *p* < 0.001 (***); ns = not significant.

**S3 Fig.**
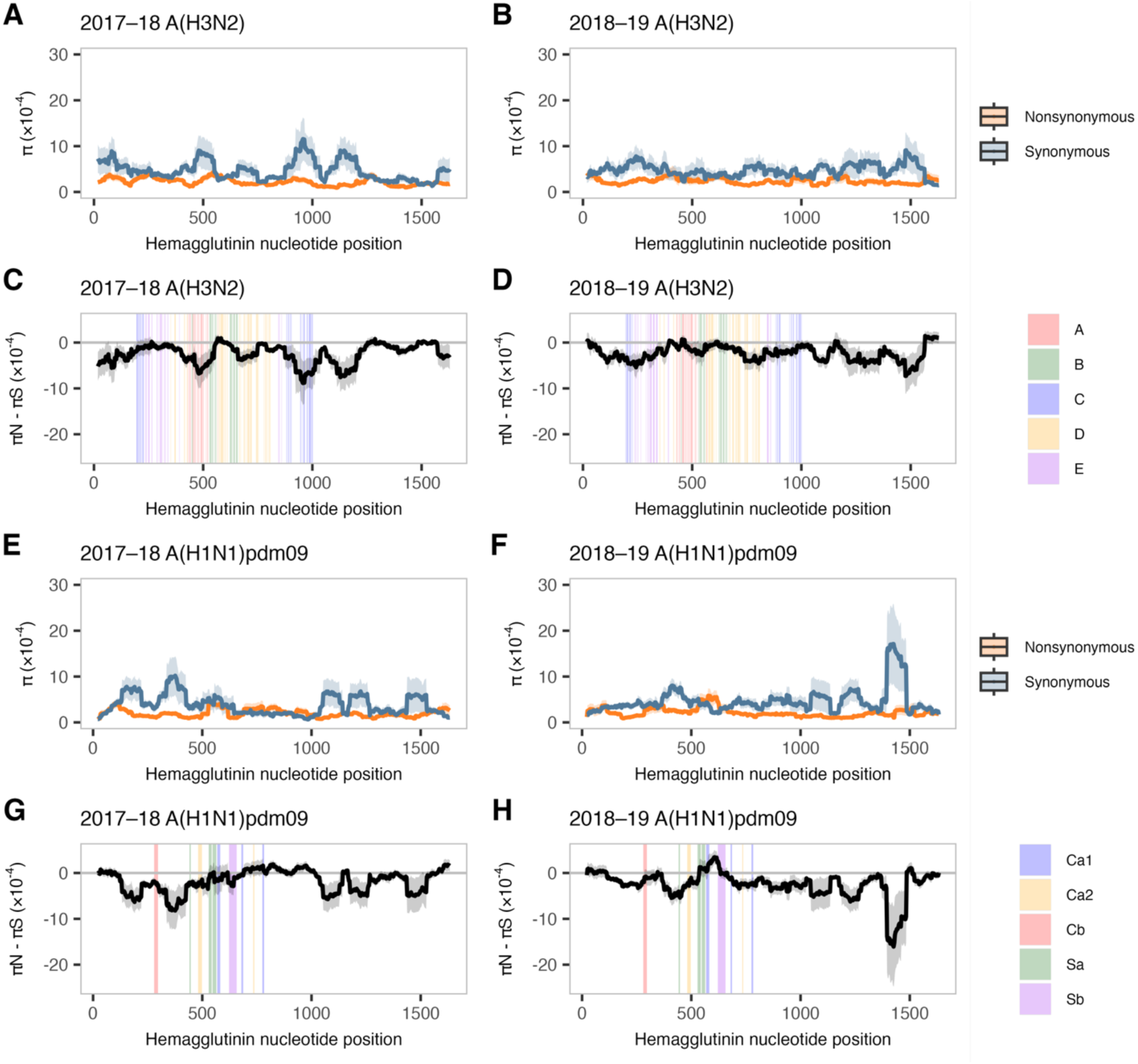
Sliding-window hemagglutinin nucleotide diversity across influenza seasons and subtypes. **(A**, **B**, **E**, **F)** Mean *π_N_* (orange) and *π_S_* (blue) values for 30-codon sliding windows in the hemagglutinin gene for **(A)** 2017–18 A(H3N2), **(B)** 2018–19 A(H3N2), **(E)** 2017–18 A(H1N1)pdm09, and **(F)** 2018–19 A(H1N1)pdm09. **(C**, **D**, **G**, **H)** Mean *π_N_* – *π_S_* values (black) for 30-codon sliding windows in the hemagglutinin gene for **(C)** 2017–18 A(H3N2), **(D)** 2018–19 A(H3N2), **(G)** 2017–18 A(H1N1)pdm09, and **(H)** 2018–19 A(H1N1)pdm09, colored by respective antigenic region. Shaded areas denote the standard error for each codon position.

**S4 Fig.**
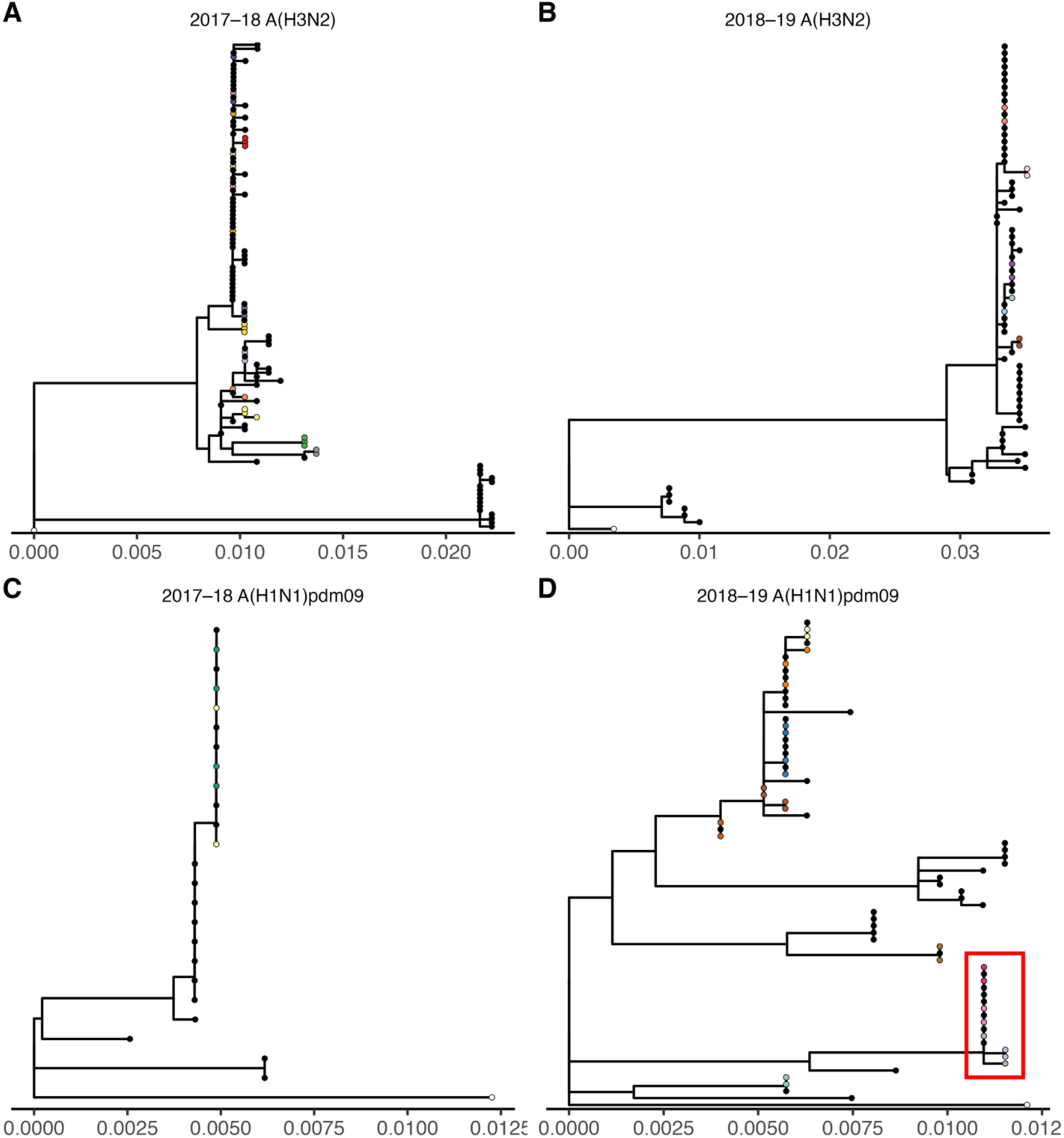
Maximum-likelihood hemagglutinin phylogenies by influenza season and subtype. **(A)** 2017–18 A(H3N2), **(B)** 2018–19 A(H3N2), **(C)** 2017–18 A(H1N1)pdm09, and **(D)** 2018–19 A(H1N1)pdm09 (divergent taxon boxed in red). Tips for validated donor–recipient pairs are colored by household; all other tips are black. The season-appropriate vaccine strain reference used as the outgroup is shown in white. Trees were inferred with RAxML-NG under GTR+Gamma and rooted to the vaccine reference; scale bars indicate substitutions per site.

**S5 Fig.**
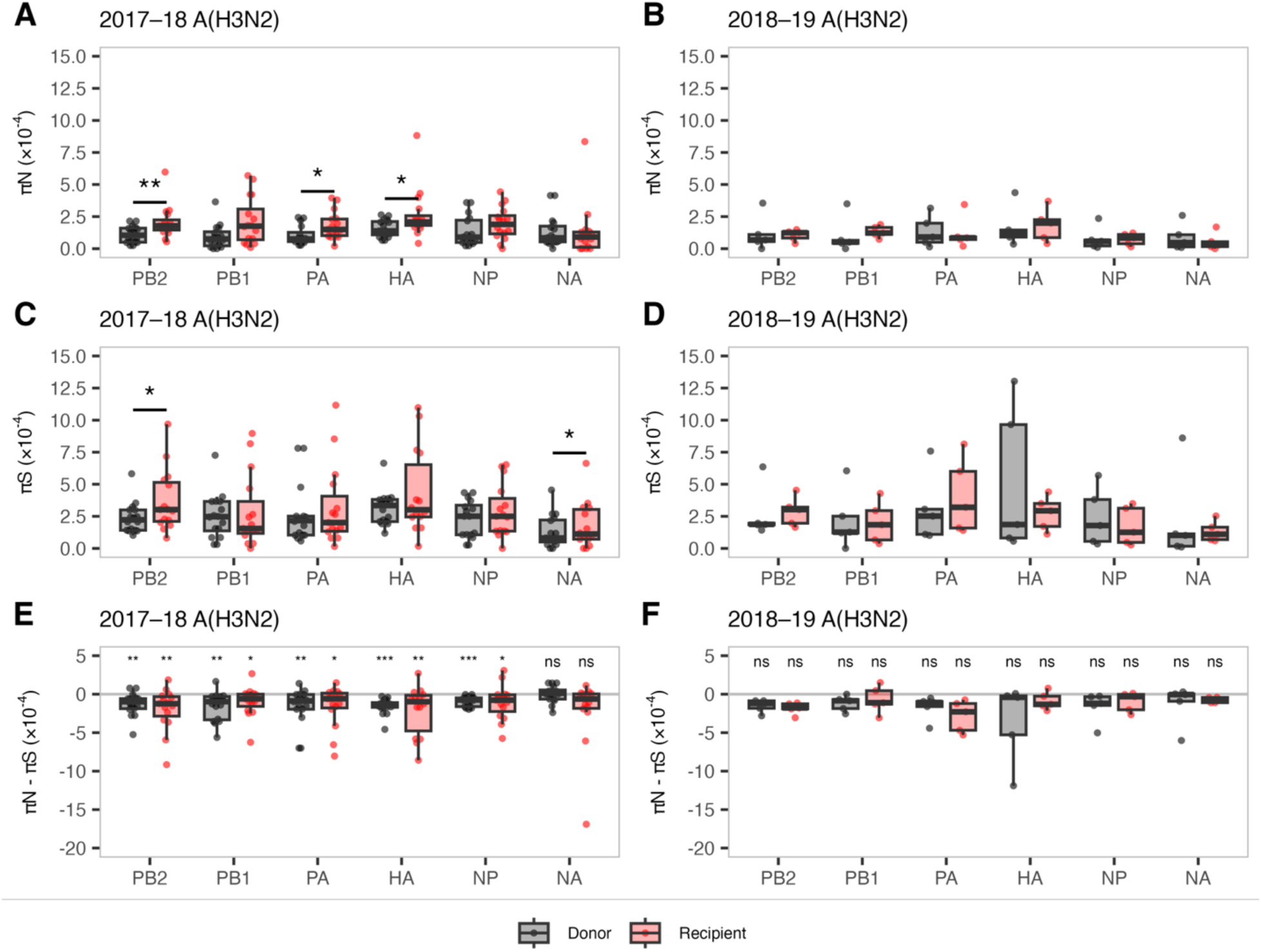
Gene-wise *π_N_*, *π_S_*, and *π_N_* – *π_S_* values in A(H3N2) donors and recipients. **(A** and **B)** *π_N_*, **(C** and **D)** *π_S_*, and **(E** and **F)** *π_N_* – *π_S_* values for all donor (black) and recipient (red) samples across all genes included in this study, for 2017–18 A(H3N2) and 2018–19 A(H3N2), respectively. Genes shown: PB2 (polymerase basic 2), PB1 (polymerase basic 1), PA (polymerase acidic), HA (hemagglutinin), NP (nucleoprotein), and NA (neuraminidase). Not included: MP (matrix; M1/M2) and NS (nonstructural; NS1/NEP) due to overlapping reading frames. The horizontal grey line in E and F represents zero, where *π_N_* = *π_S_*. Paired comparisons of *π_N_* and *π_S_* between donors and recipients using Wilcoxon signed-rank tests. Median differences between *π_N_* and *π_S_* were tested in against zero using one-sample Wilcoxon signed-rank tests. Asterisks indicate significance thresholds: *p* < 0.05 (*), *p* < 0.01 (**), and *p* < 0.001 (***); ns = not significant.

**S6 Fig.**
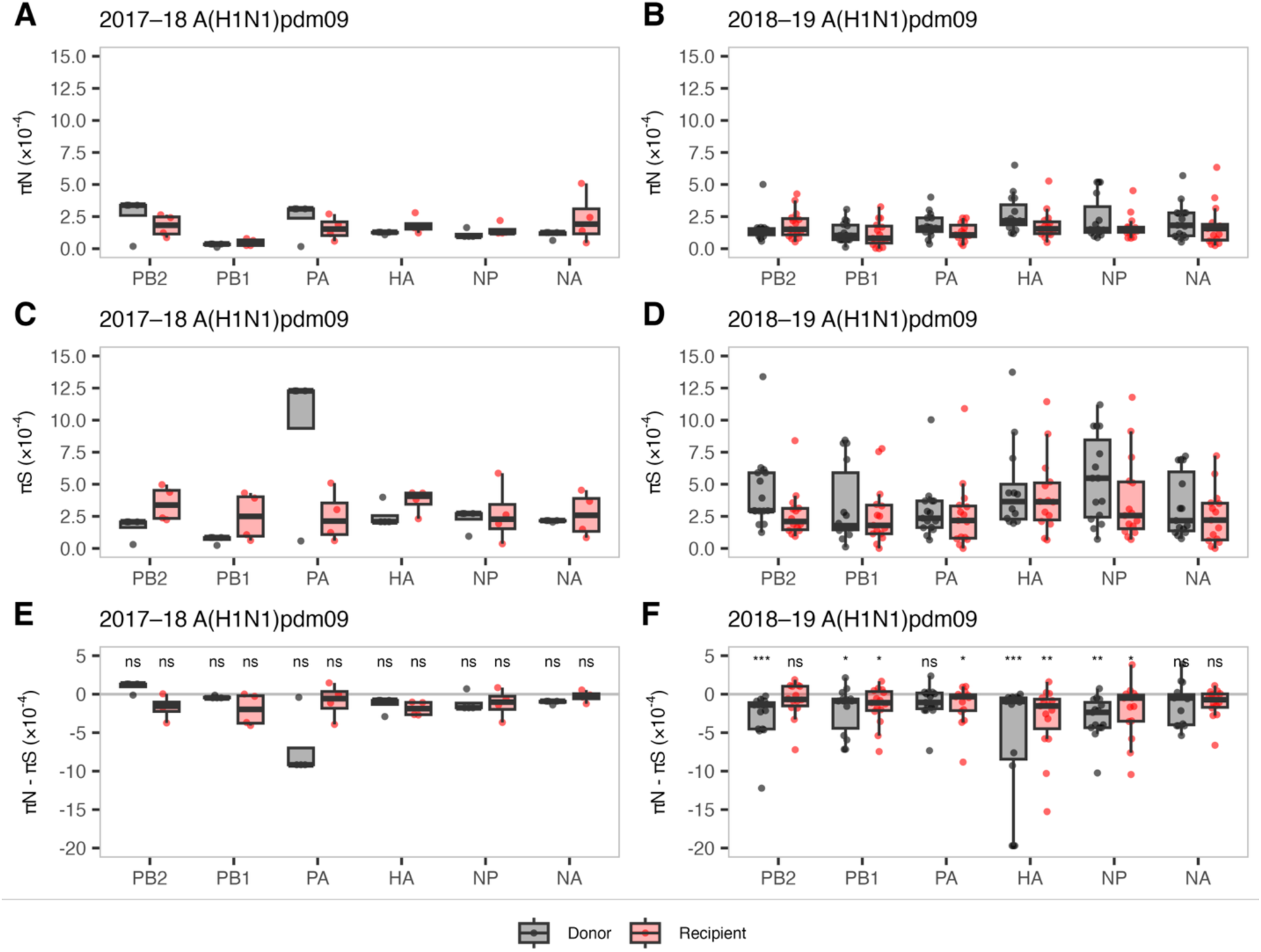
Gene-wise *π_N_*, *π_S_*, and *π_N_* – *π_S_* values in A(H1N1)pdm09 donors and recipients. **(A** and **B)** *π_N_*, **(C** and **D)** *π_S_*, and **(E** and **F)** *π_N_* – *π_S_* values for all donor (black) and recipient (red) samples across all genes included in this study, for 2017–18 A(H1N1)pdm09 and 2018–19 A(H1N1)pdm09, respectively. Genes shown: PB2 (polymerase basic 2), PB1 (polymerase basic 1), PA (polymerase acidic), HA (hemagglutinin), NP (nucleoprotein), and NA (neuraminidase). Not included: MP (matrix; M1/M2) and NS (nonstructural; NS1/NEP) due to overlapping reading frames. The horizontal grey line in E and F represents 0, where *π_N_* = *π_S_*. Paired comparisons of *π_N_* and *π_S_* between donors and recipients using Wilcoxon signed-rank tests. Median differences between *π_N_* and *π_S_* were tested in against zero using one-sample Wilcoxon signed-rank tests. Asterisks indicate significance thresholds: *p* < 0.05 (*), *p* < 0.01 (**), and *p* < 0.001 (***); ns = not significant.

**S7 Fig.**
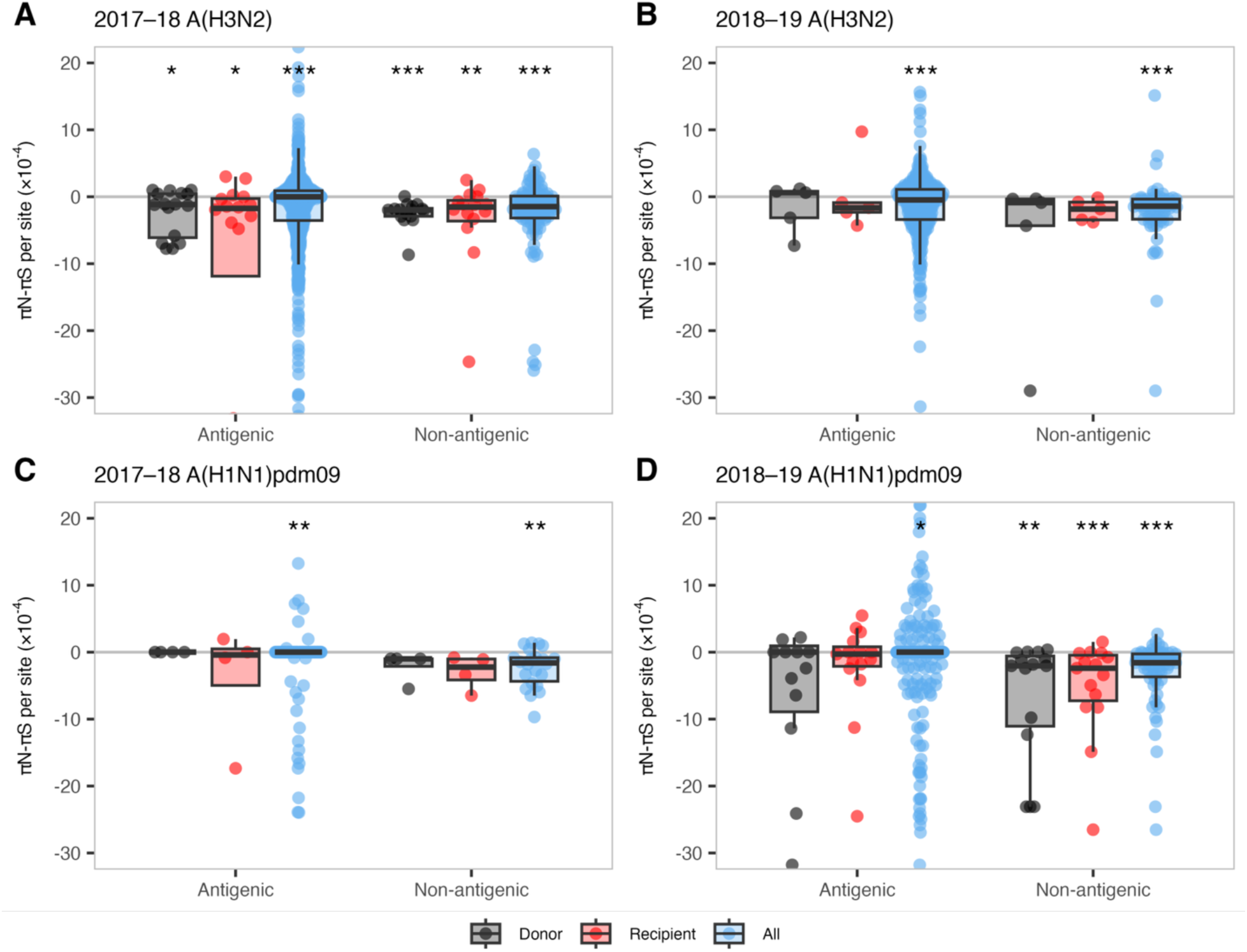
Per-site *π_N_* – *π_S_* values in antigenic and non-antigenic regions of HA. **(A)** 2017–18 A(H3N2), **(B)** 2018–19 A(H3N2), **(C)** 2017–18 A(H1N1)pdm09, and **(D)** 2018–19 A(H1N1)pdm09. *π_N_* – *π_S_* values were normalized to account for differences in the amount of antigenic and non-antigenic sites, denoted *π_N_* – *π_S_* per site. Boxplots are overlaid to indicate the median, interquartile range, and values within 1.5× the interquartile range for each group: Donor (black), Recipient (red), or All (blue). The “All” group includes values for all viral genomes sampled (donors, recipients, and also those without a linked household contact). The horizontal grey line represents zero, where *π_N_* = *π_S_*. Median differences between *π_N_* and *π_S_* were tested against zero using one-sample Wilcoxon signed-rank tests. Asterisks indicate significance thresholds: *p* < 0.05 (*), *p* < 0.01 (**), and *p* < 0.001 (***); ns = not significant.

**S8 Fig.**
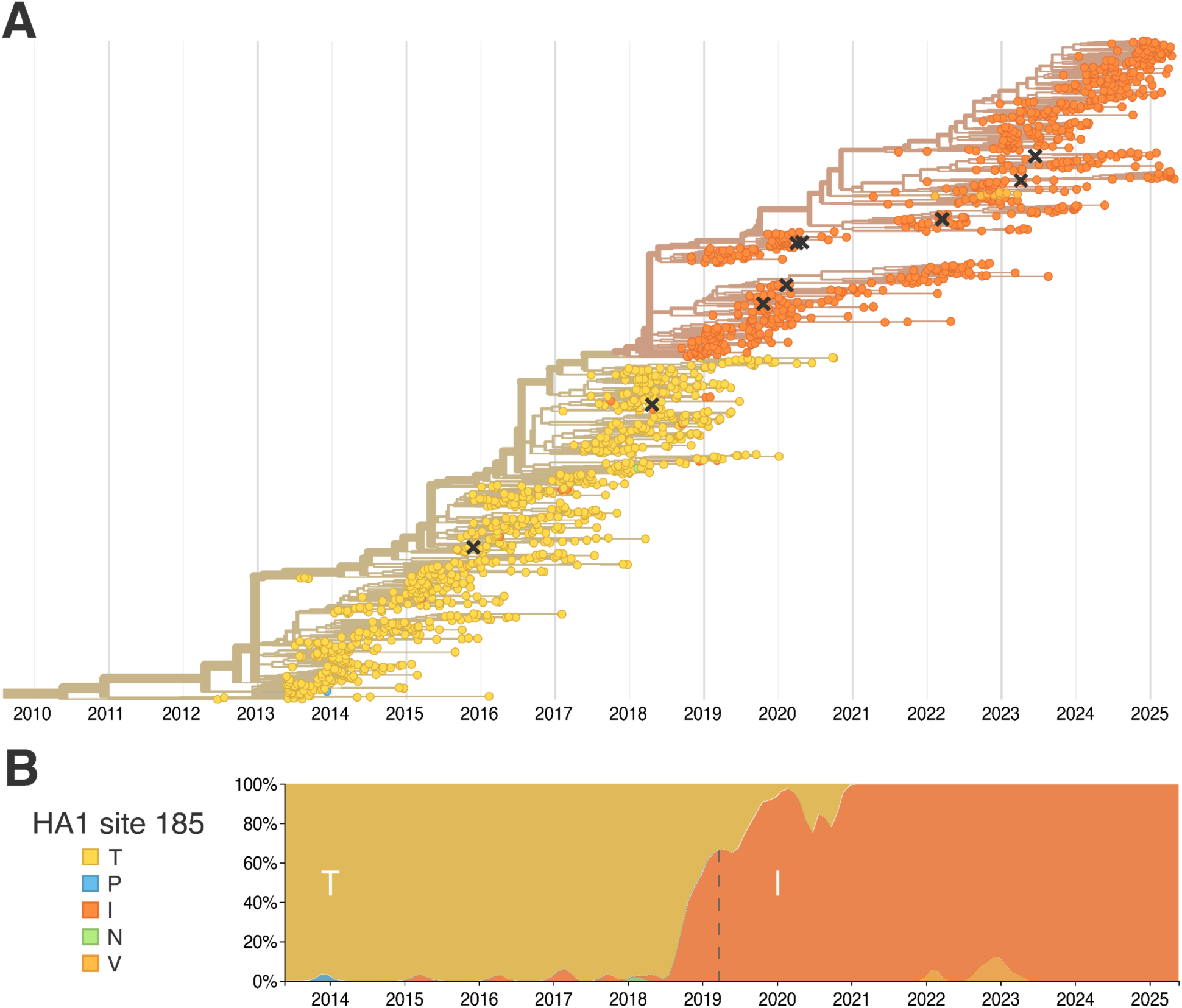
Global dynamics of HA1 site 185 amino acid identities. **(A)** Time-resolved phylogeny of H1 consensus sequences (n = 1,461), constructed using Nextstrain and GISAID data. Branches and tips are colored by amino acid identity at HA1 site 185: threonine (T, yellow), proline (P, blue), isoleucine (I, dark orange), asparagine (N, green), and valine (V, light orange). **(B)** Temporal dynamics of HA1 185 genotypes among global sequences, showing the proportion of each amino acid over time. The dashed vertical line marks the approximate timing of I185T fixation in a specific donor-recipient pair, during a period when 185I was present in 66% of global sequences. Both panels were adapted from Nextstrain [41] and modified to align x-axes and improve label clarity.

**S9 Fig.**
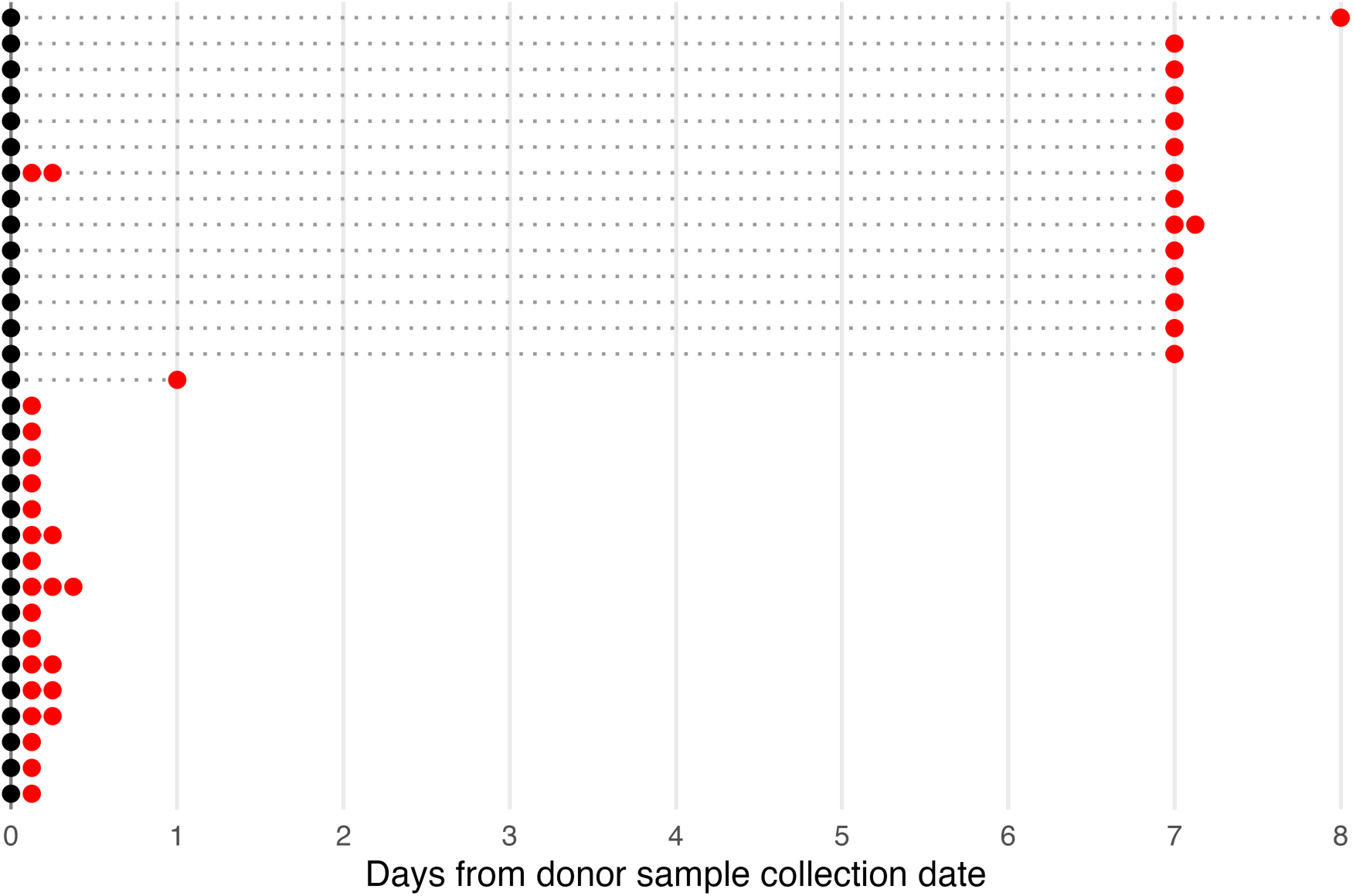
Household sample collection timelines relative to donor sample collection date. Sample collection timelines for the 31 households meeting both epidemiologic and genetic criteria for influenza transmission. Each row represents a single household, with the donor sample shown as a black circle and recipient sample(s) shown as red circles. Dotted gray lines connect each donor to their corresponding recipient(s). When multiple samples within a household were collected on the same day, recipient points are offset by +0.125 days along the x-axis to avoid overlap.

**S1 Table.**
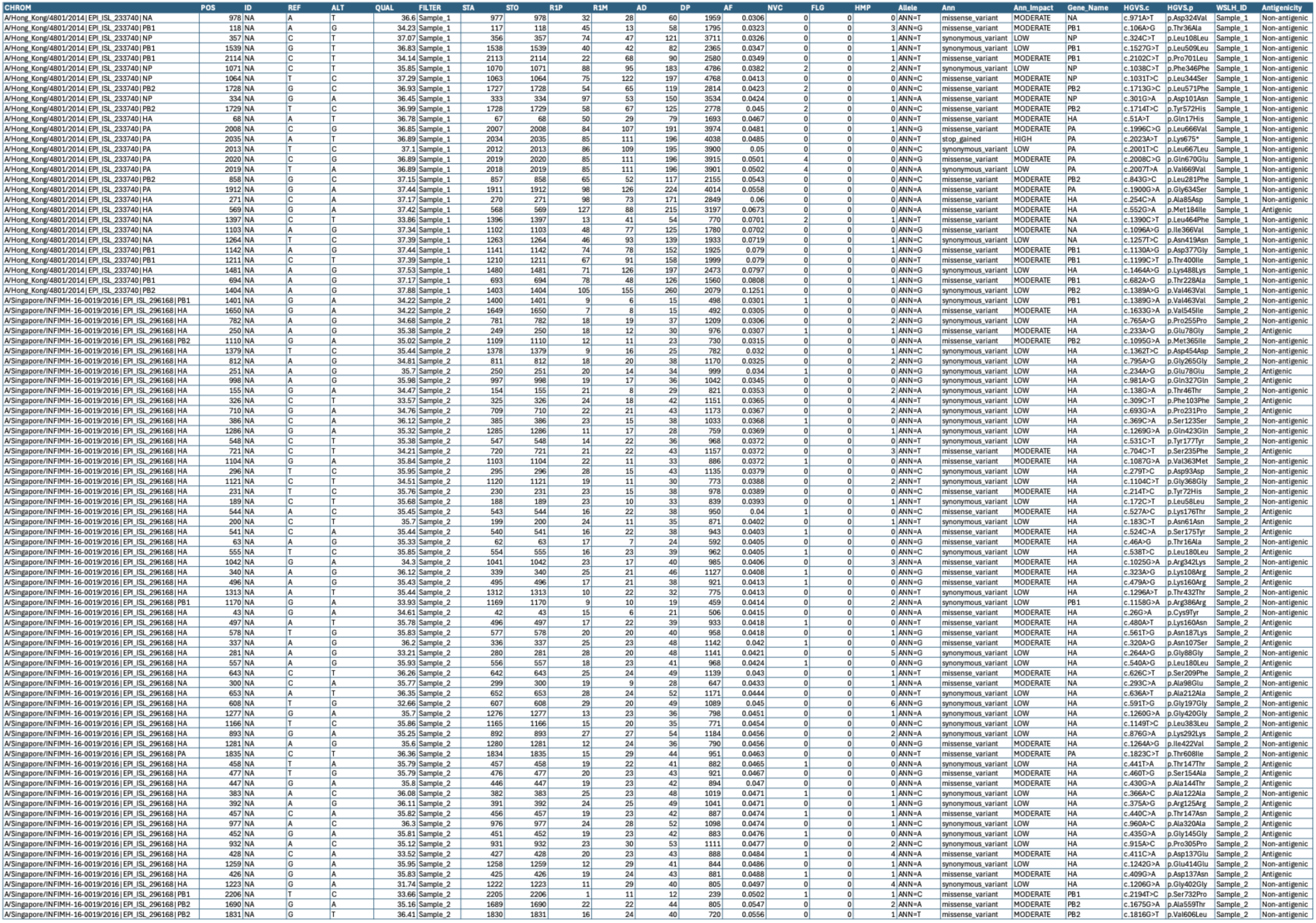
Within-host variants from two outlier samples. Intra-host single-nucleotide variants (iSNVs) from two samples with greater than twenty iSNVs. The table was generated from a Variant Call Format file from each sample. Individual sample iSNVs were annotated with snpEff, and antigenicity was annotated with RStudio (see Methods).

**S2 Table.** Notable iSNVs present in both donor and recipient samples. Merge_ID denotes the year and subtype of the sample. Genes shown: PB2 (polymerase basic 2), PB1 (polymerase basic 1), PA (polymerase acidic), HA (hemagglutinin), NP (nucleoprotein), and NA (neuraminidase). Not included: MP (matrix; M1/M2) and NS (nonstructural; NS1/NEP) due to overlapping reading frames.

| AA | iSNV | Donor_ID | Recipient_ID | Donor_AF | Recipient_AF | Merge_ID | Ann |
| --- | --- | --- | --- | --- | --- | --- | --- |
| G187G | A594G-NP | Donor 1 | Recipient 1a | 0.1003 | 0.1488 | 18-19_H1N1 | synonymous_variant |
| T22A | A97G-NP | Donor 1 | Recipient 1b | 0.242 | 0.2977 | 18-19_H1N1 | missense_variant |
| C471C | C1485T-HA | Donor 1 | Recipient 1a | 0.3164 | 0.1427 | 18-19_H1N1 | synonymous_variant |
| C471C | C1485T-HA | Donor 1 | Recipient 1b | 0.3164 | 0.7063 | 18-19_H1N1 | synonymous_variant |
| C471C | C1485T-HA | Donor 1 | Recipient 1c | 0.3164 | 0.9991 | 18-19_H1N1 | synonymous_variant |
| P627P | C1893T-PB1 | Donor 2 | Recipient 2 | 0.1045 | 0.2876 | 17-18_H3N2 | synonymous_variant |
| A270S | G823T-PB2 | Donor 3 | Recipient 3a | 0.6047 | 0.1471 | 17-18_H1N1 | missense_variant |
| A270S | G823T-PB2 | Donor 3 | Recipient 3b | 0.6047 | 0.058 | 17-18_H1N1 | missense_variant |
| N346N | T1050C-PB1 | Donor 4 | Recipient 4 | 0.5705 | 0.761 | 18-19_H1N1 | synonymous_variant |
| I185T | T626C-HA | Donor 5 | Recipient 5 | 0.3767 | 1 | 18-19_H1N1 | missense_variant |

**S3 Table.**
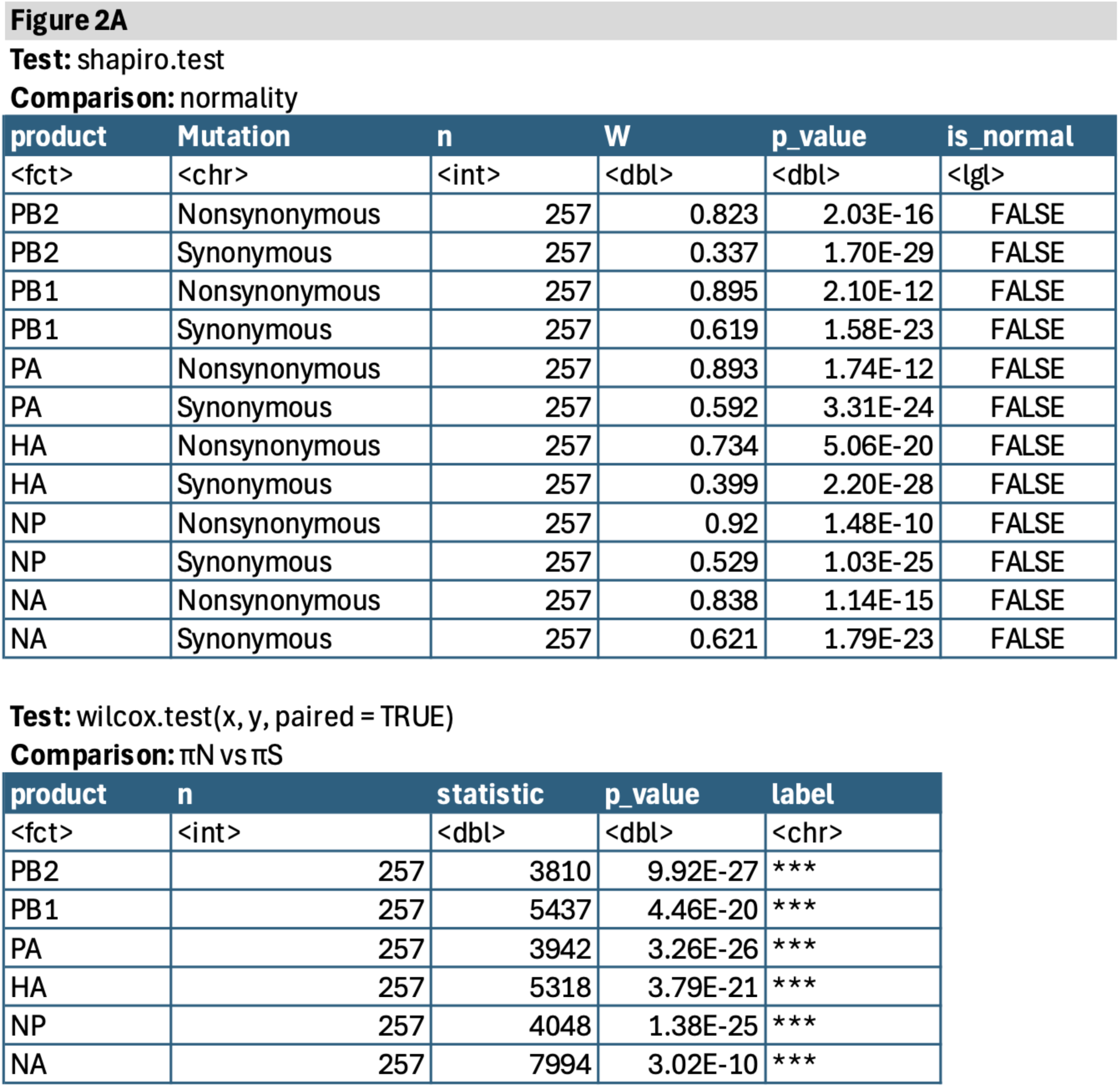
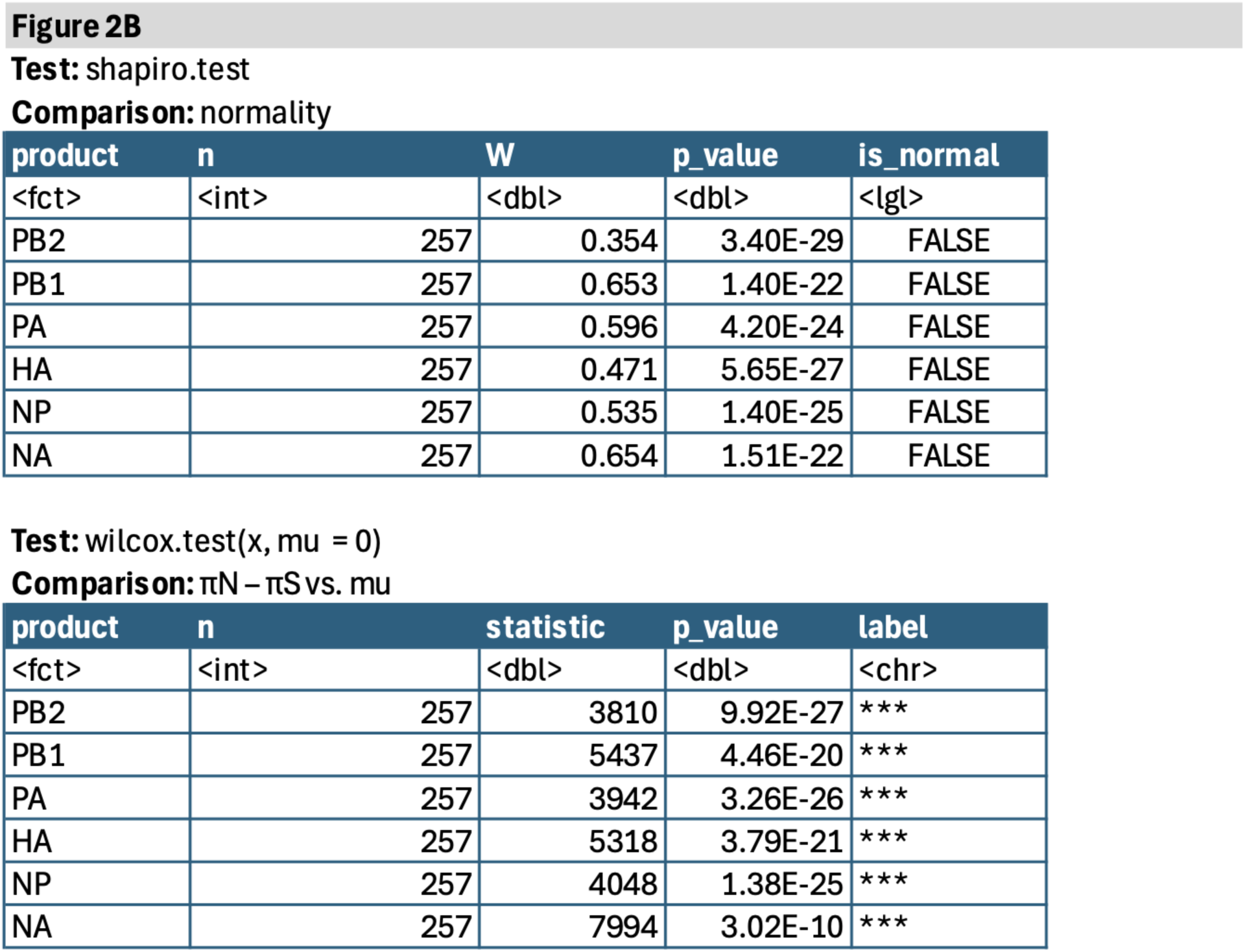
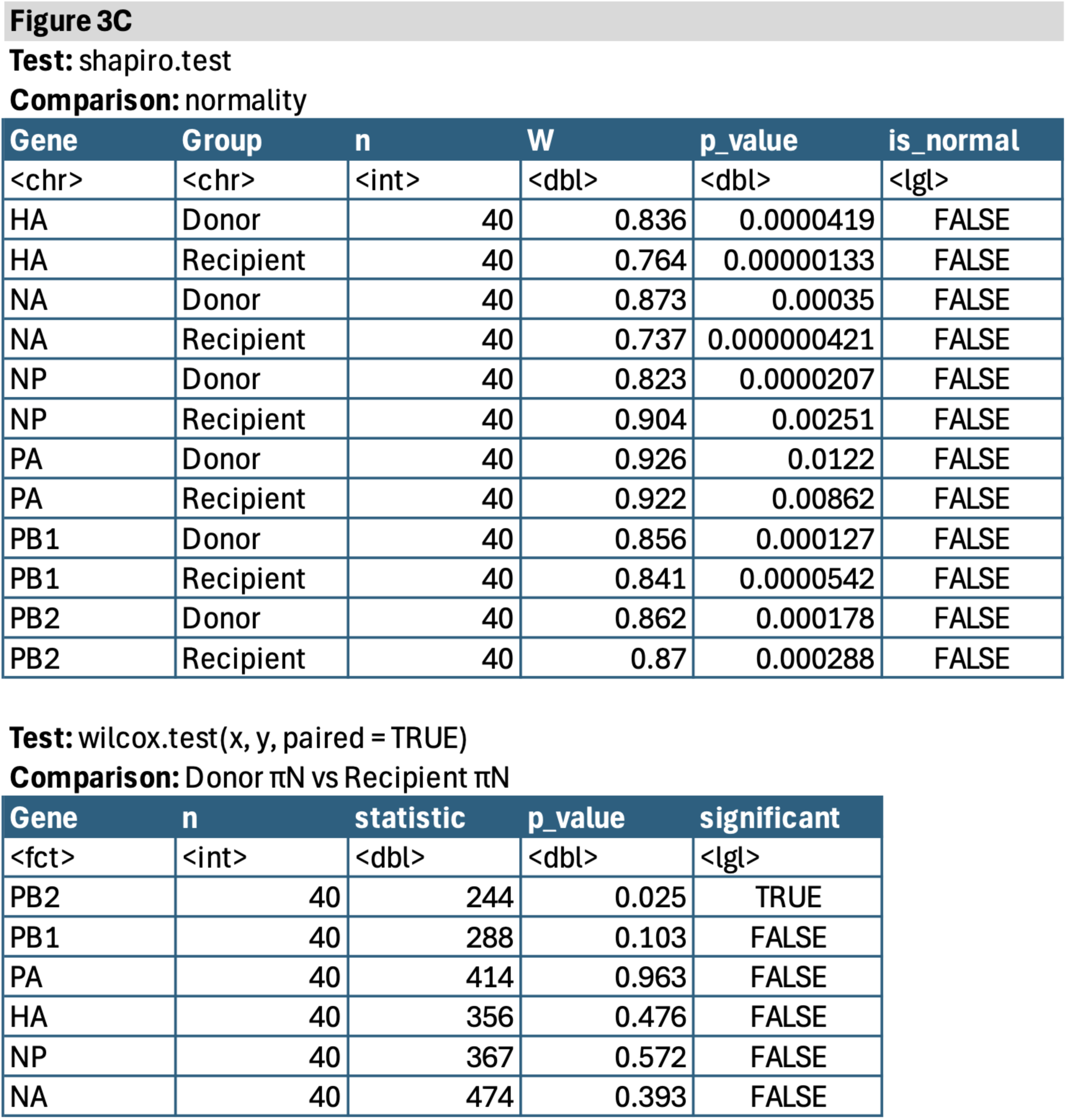
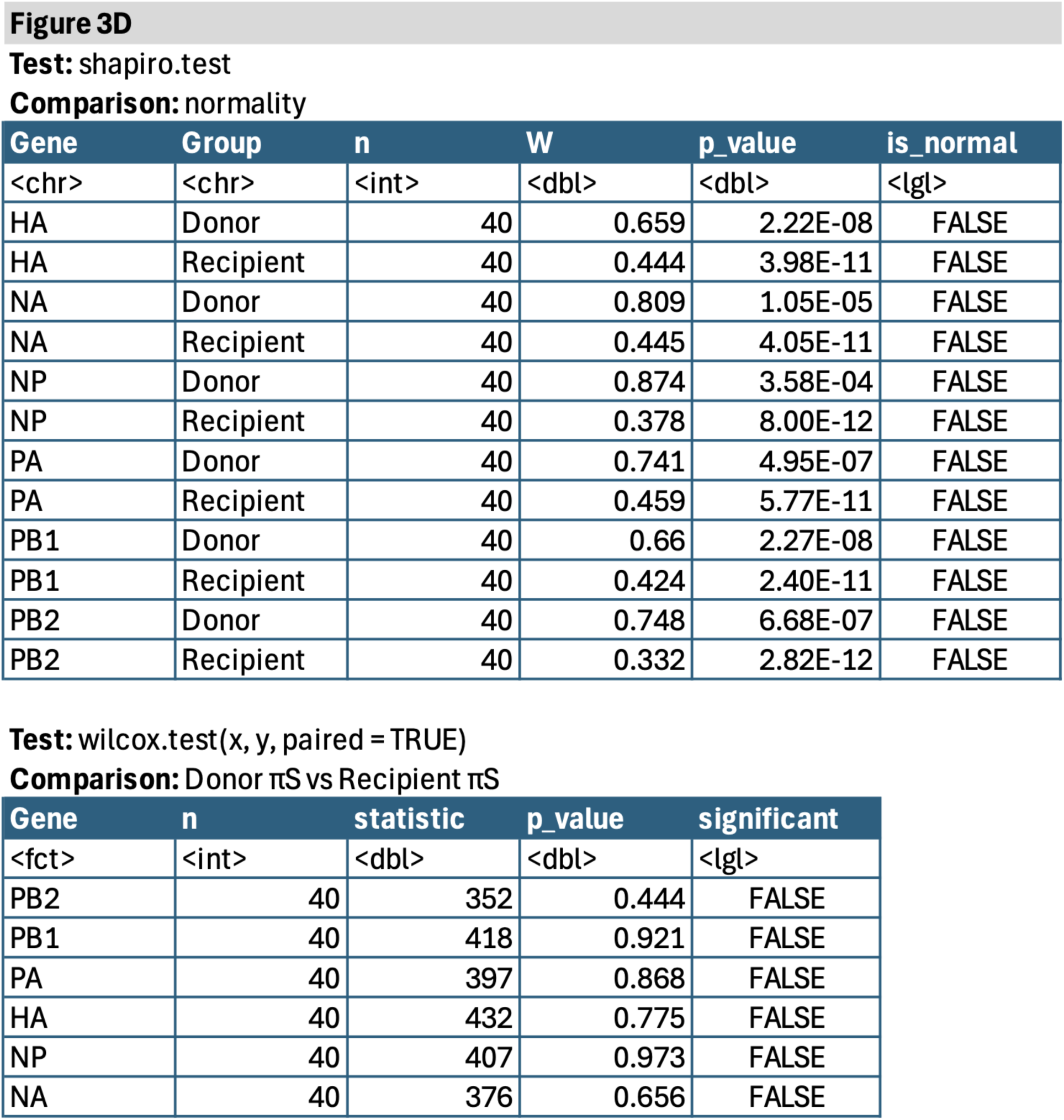
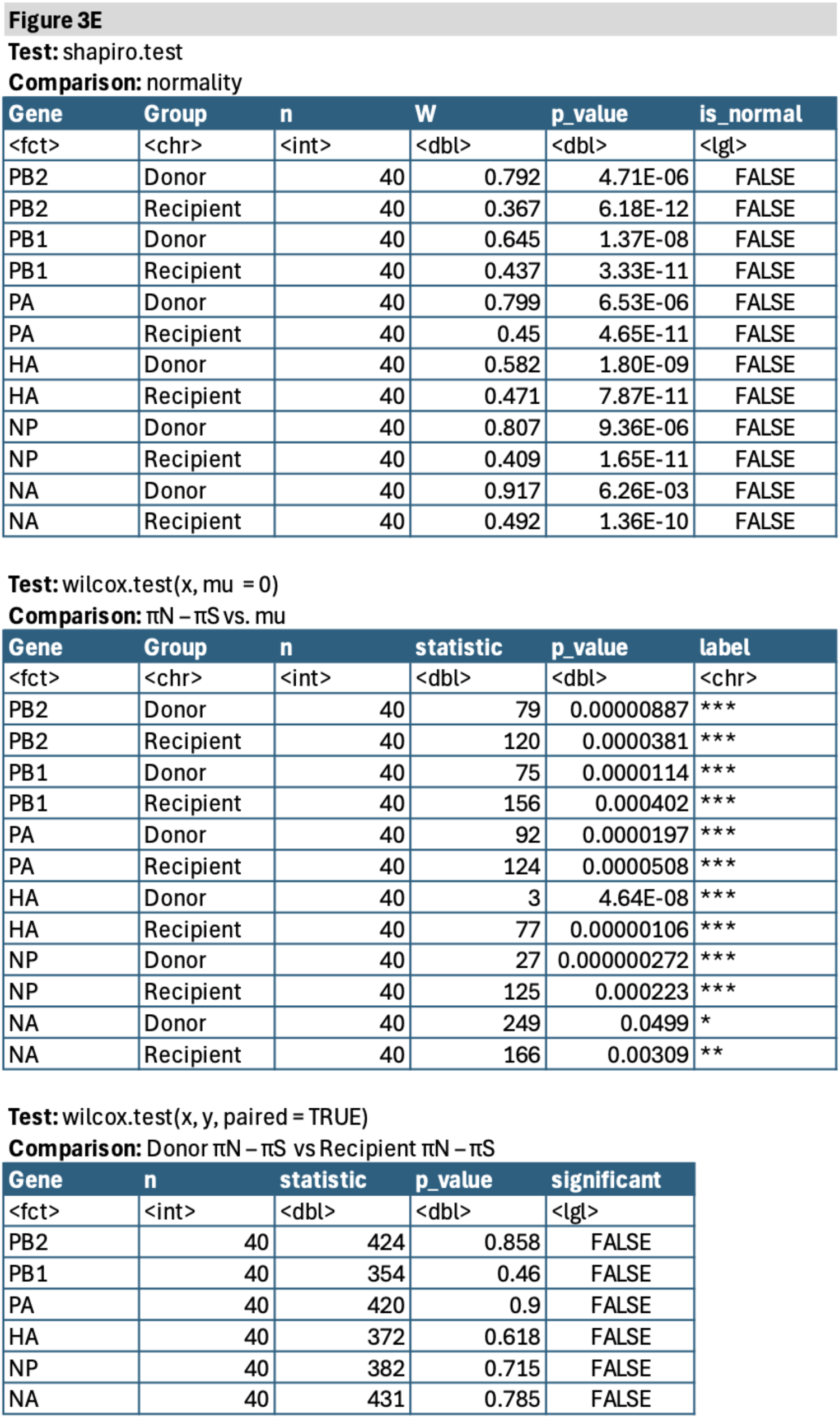
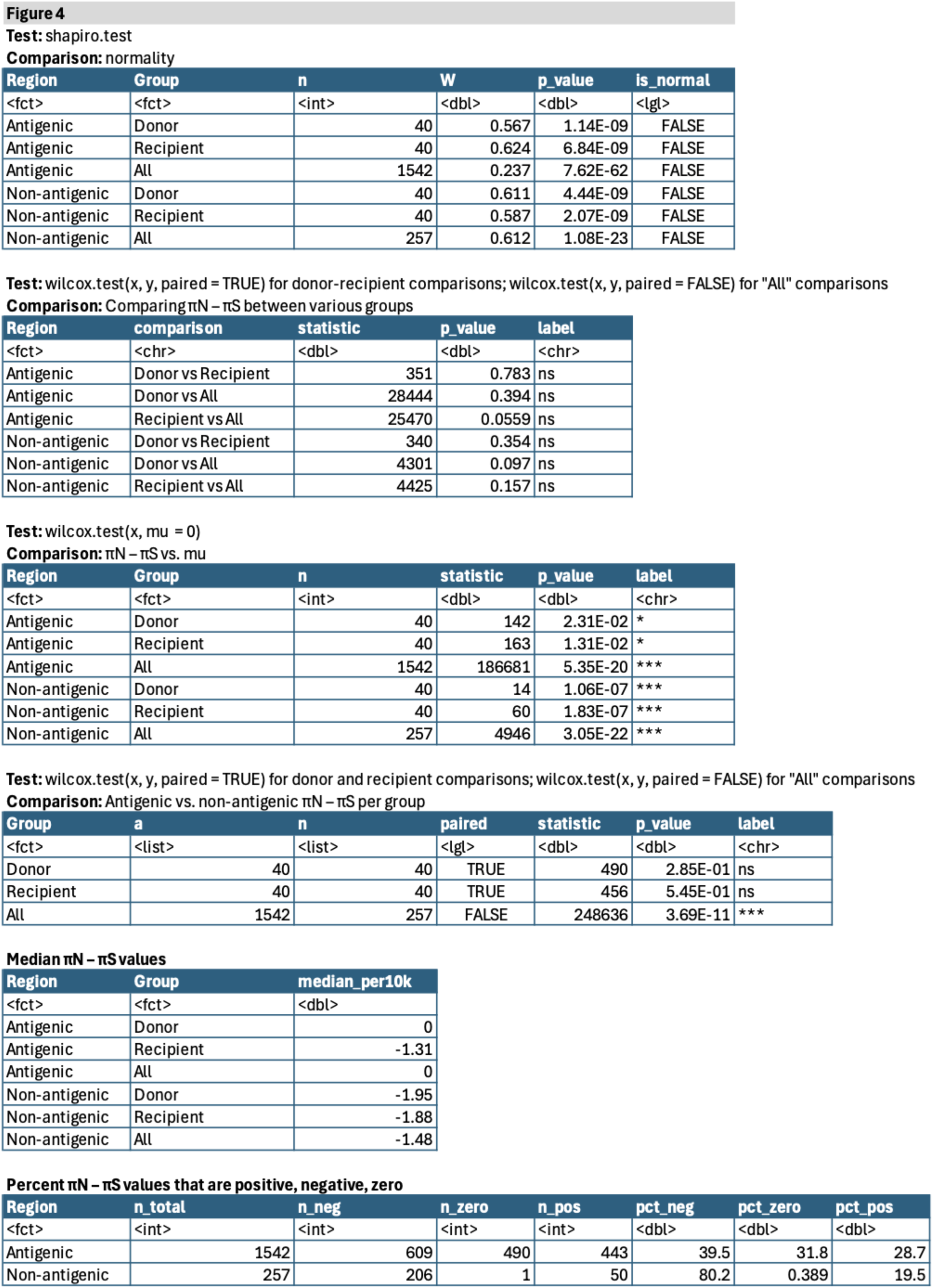
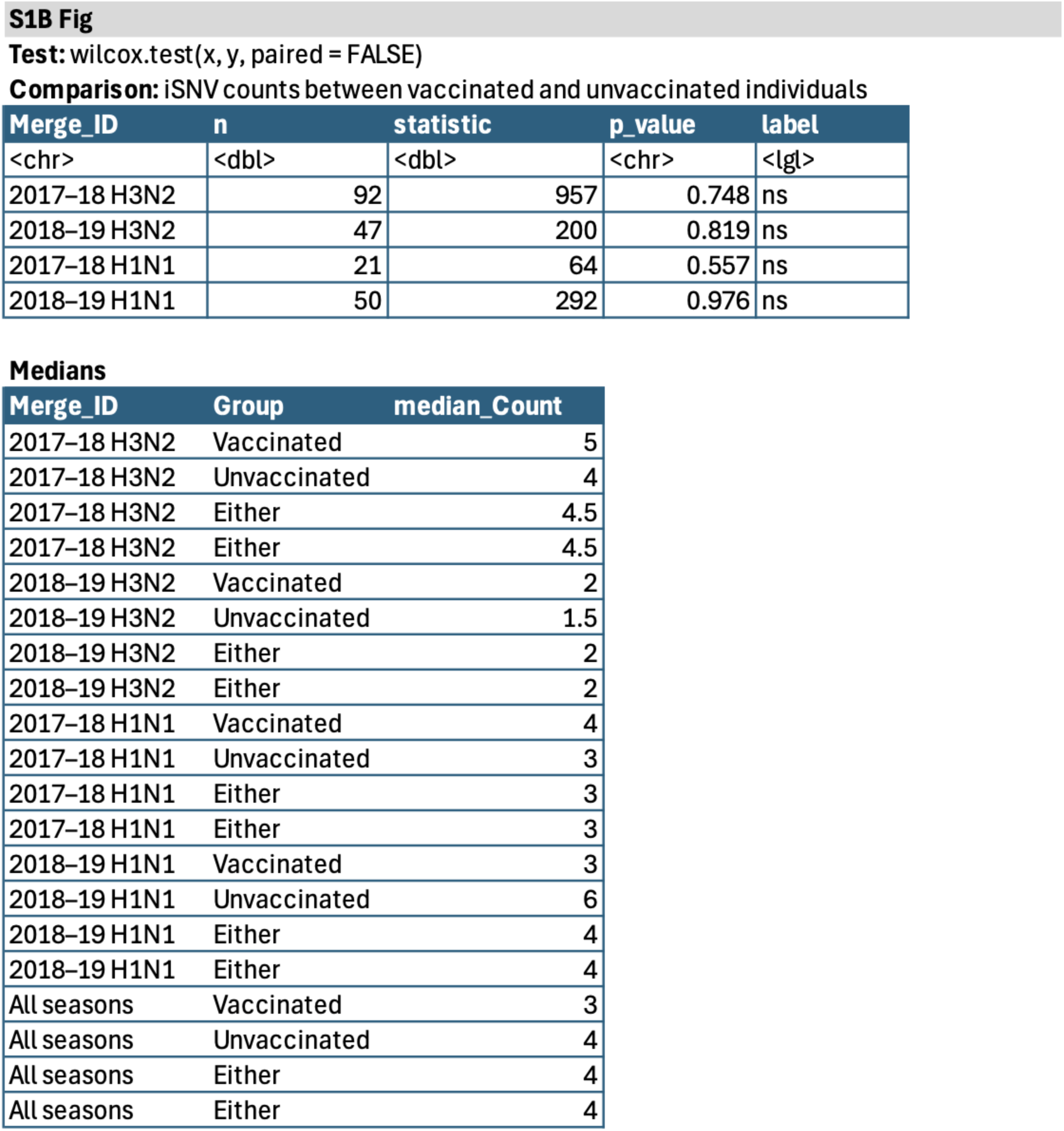
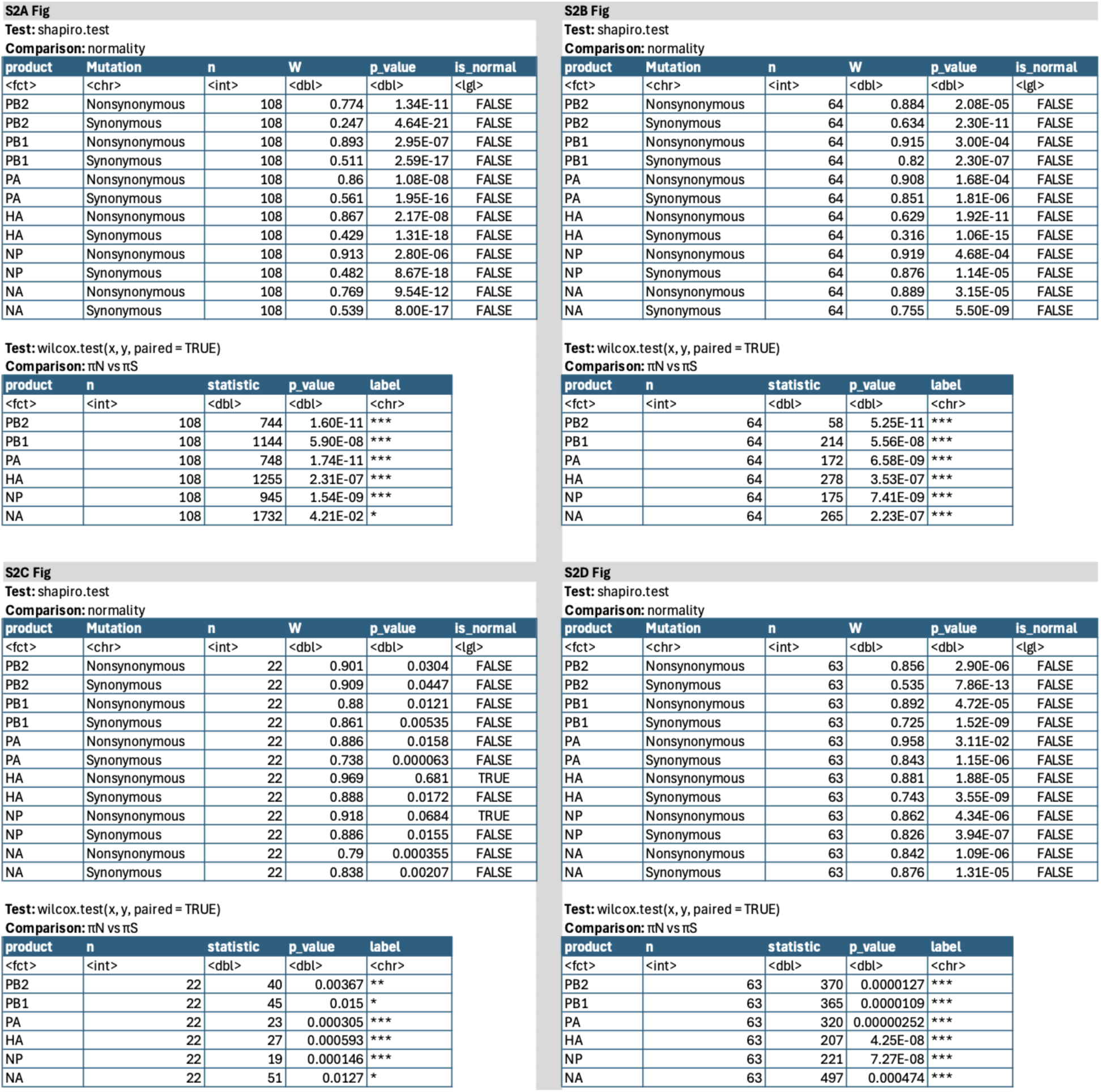
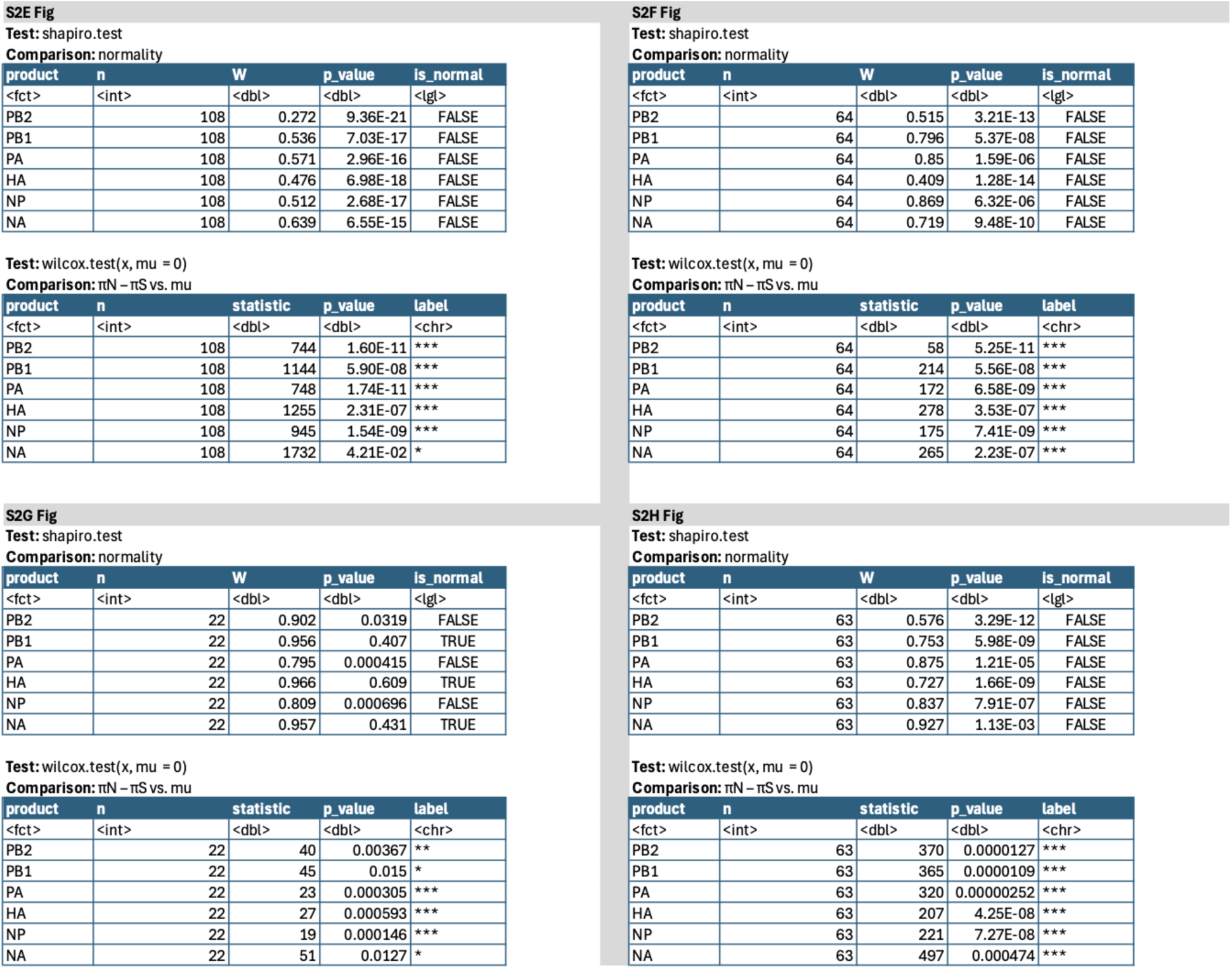
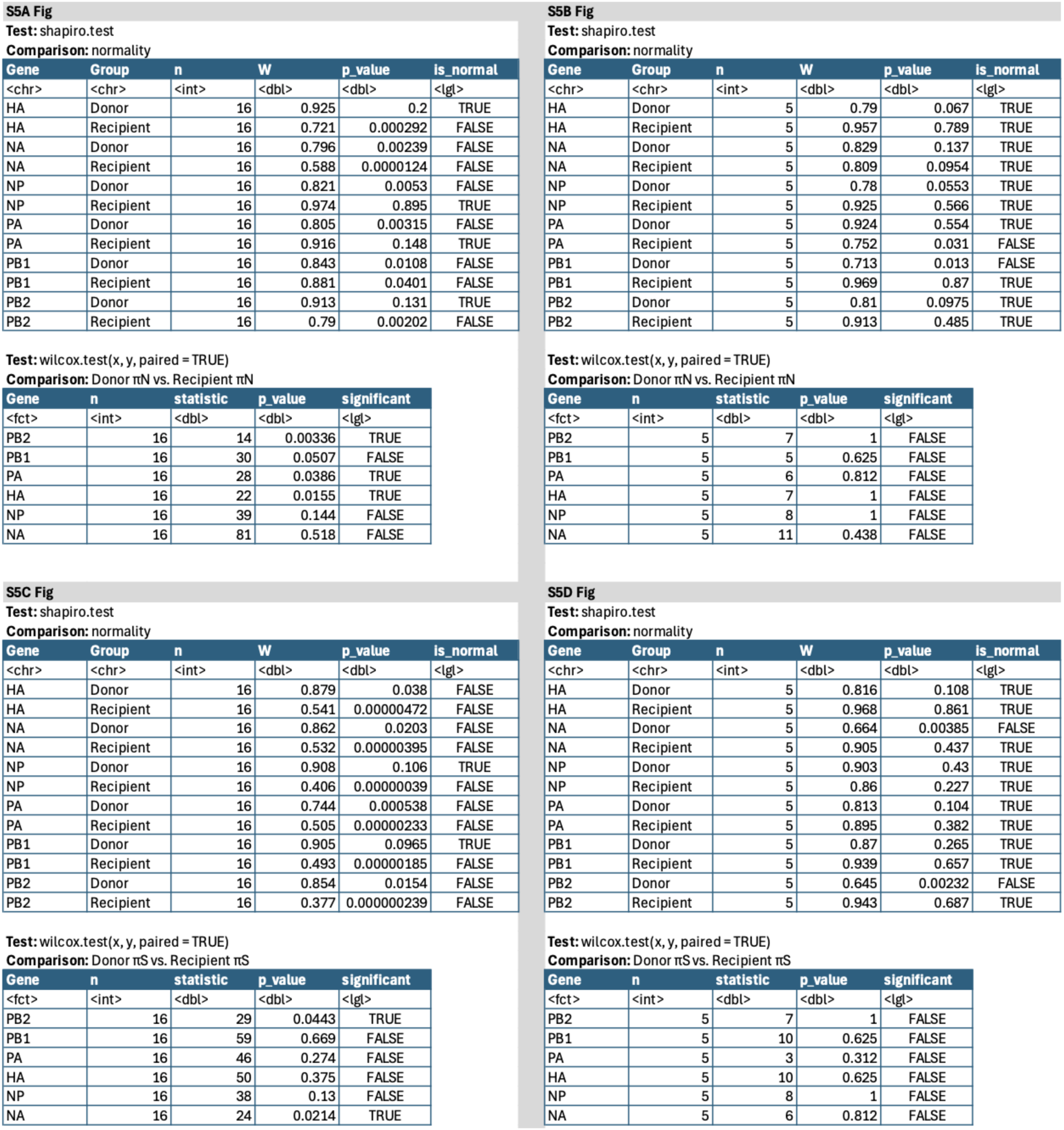
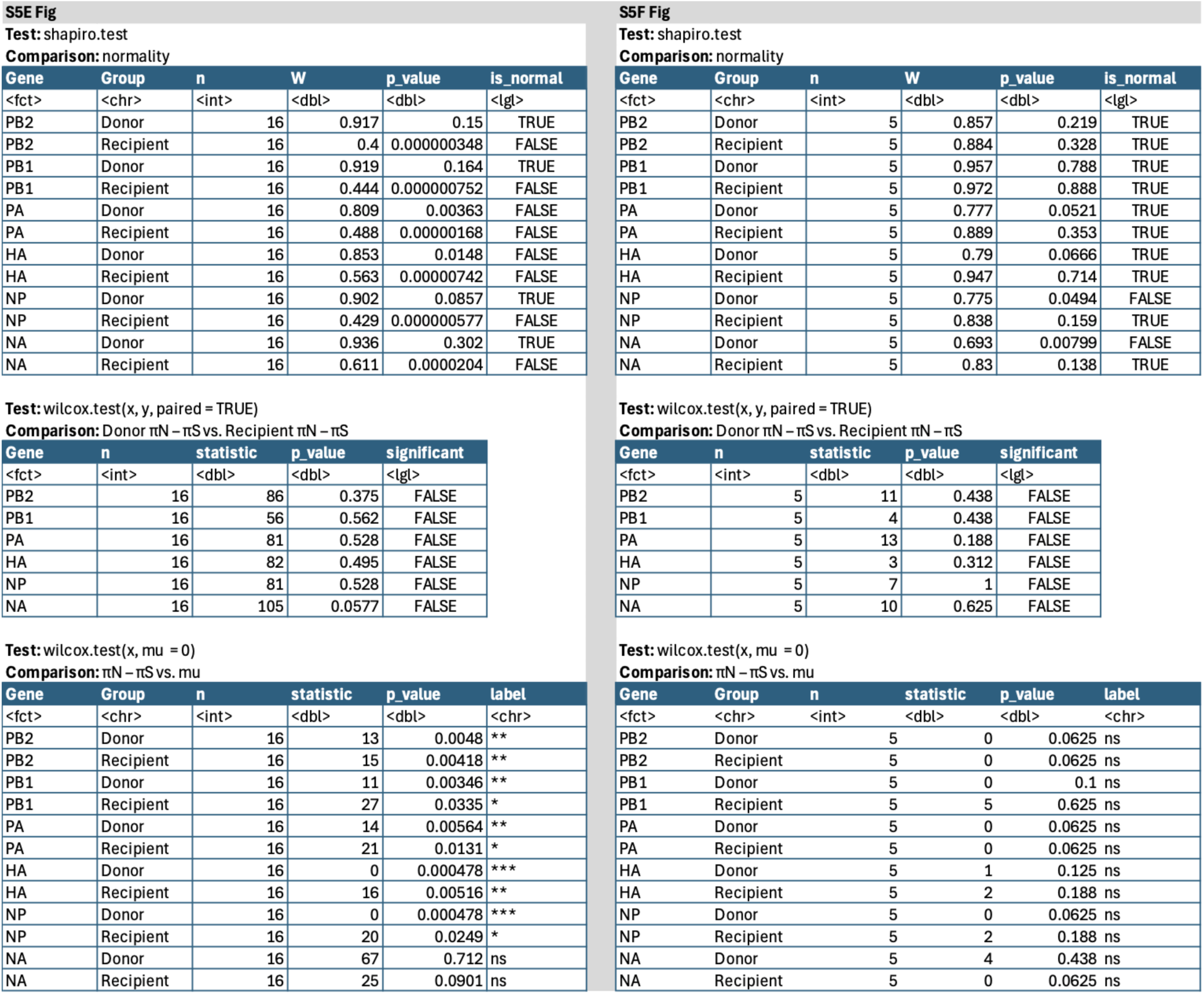
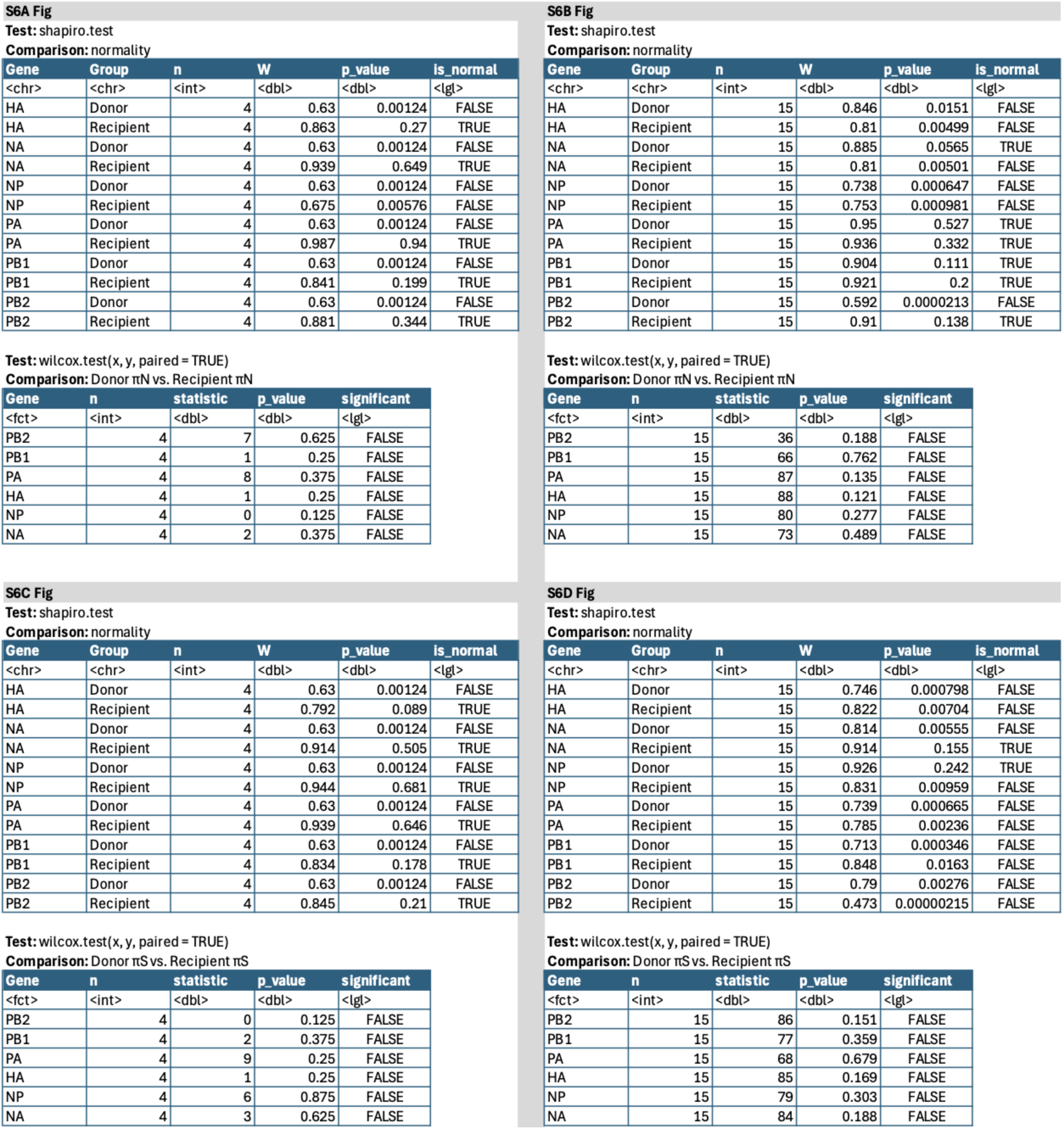
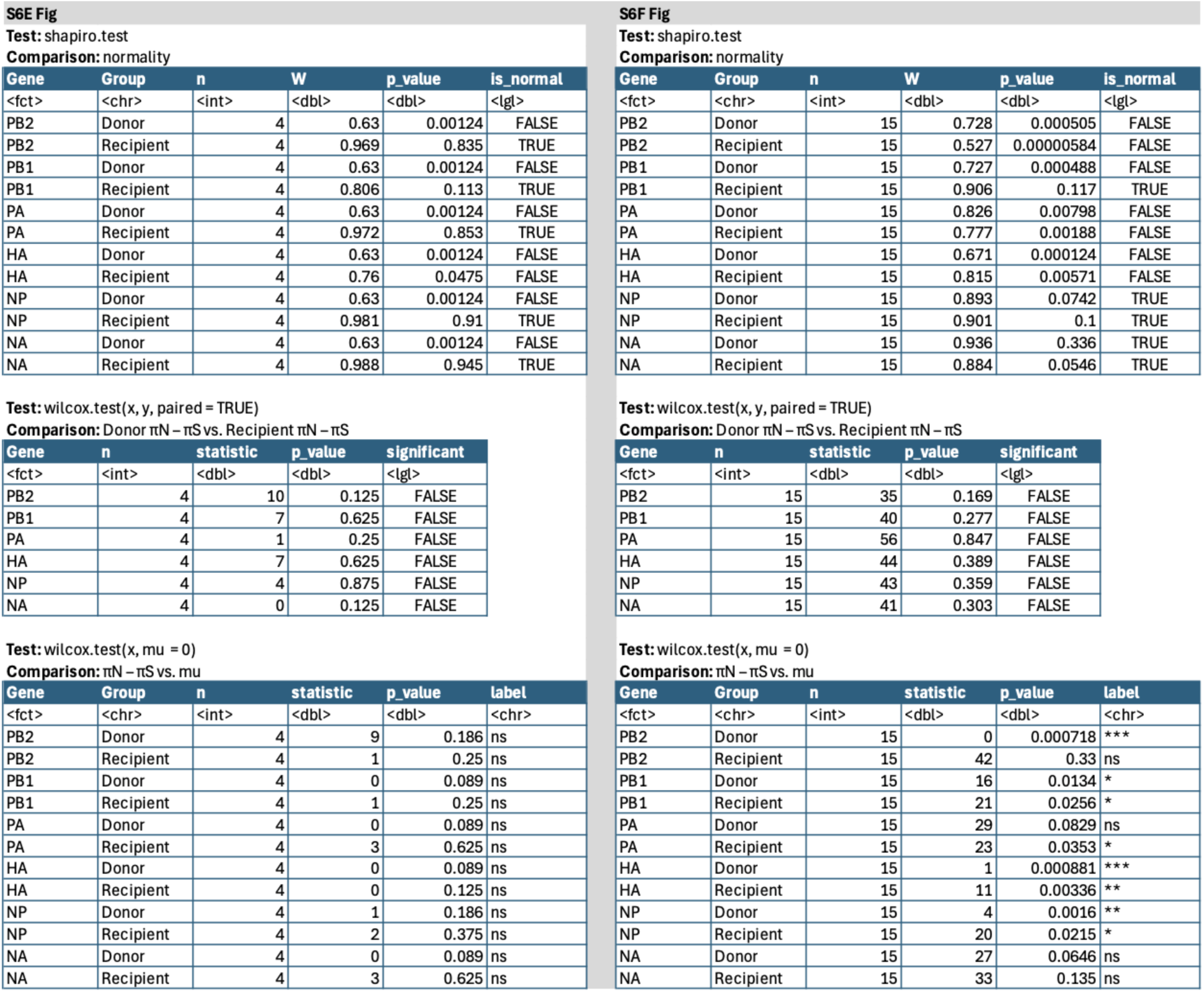
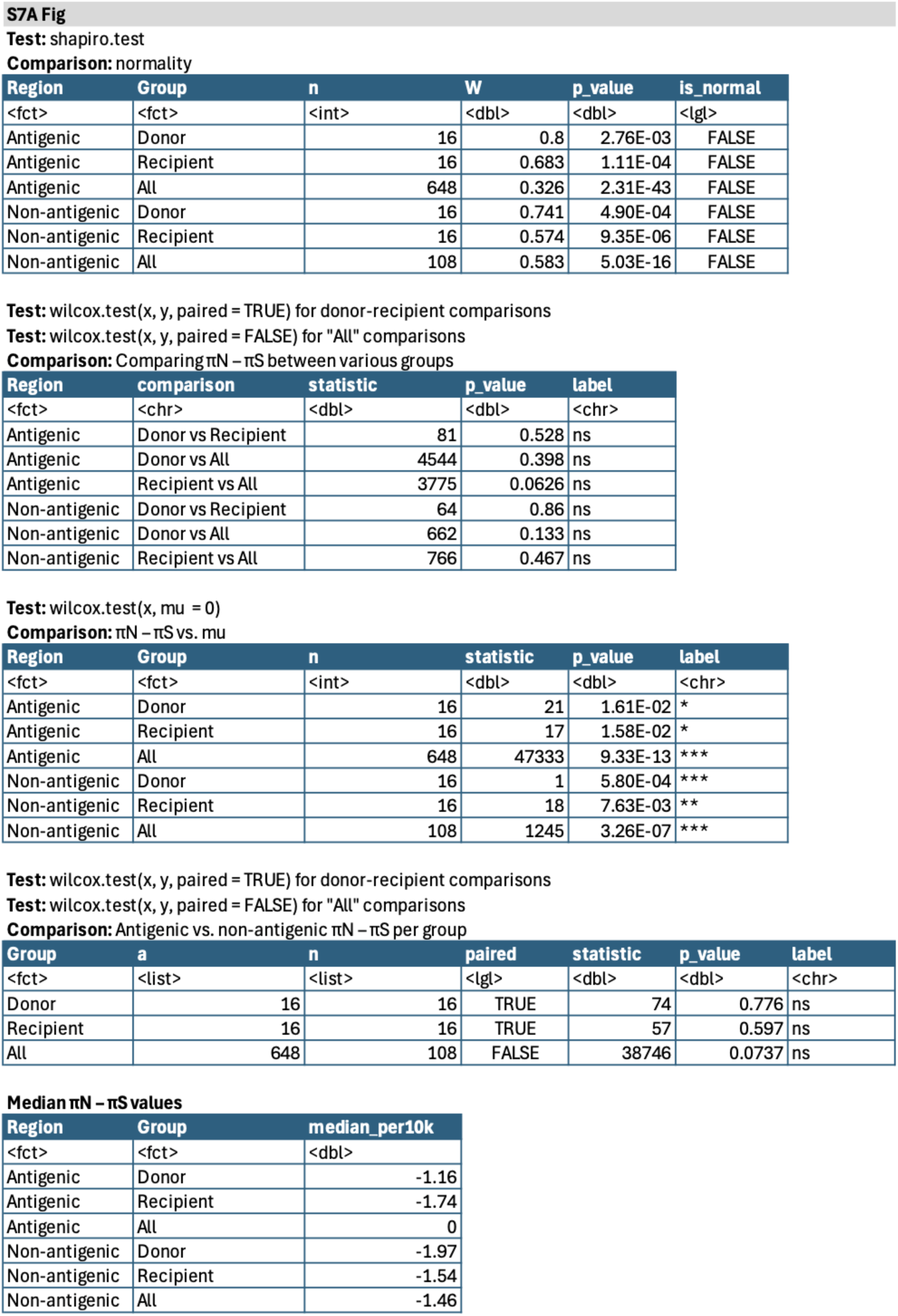
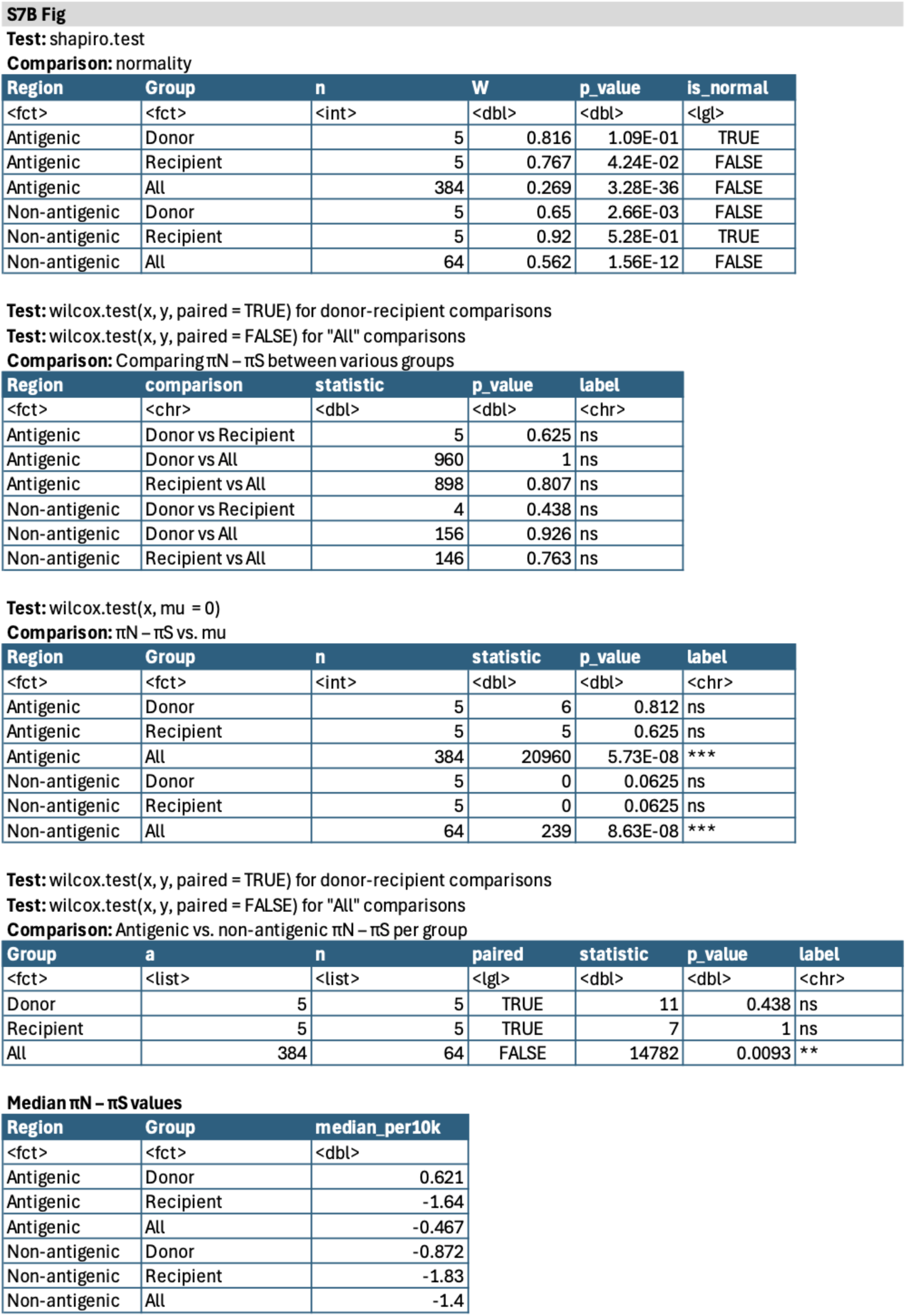
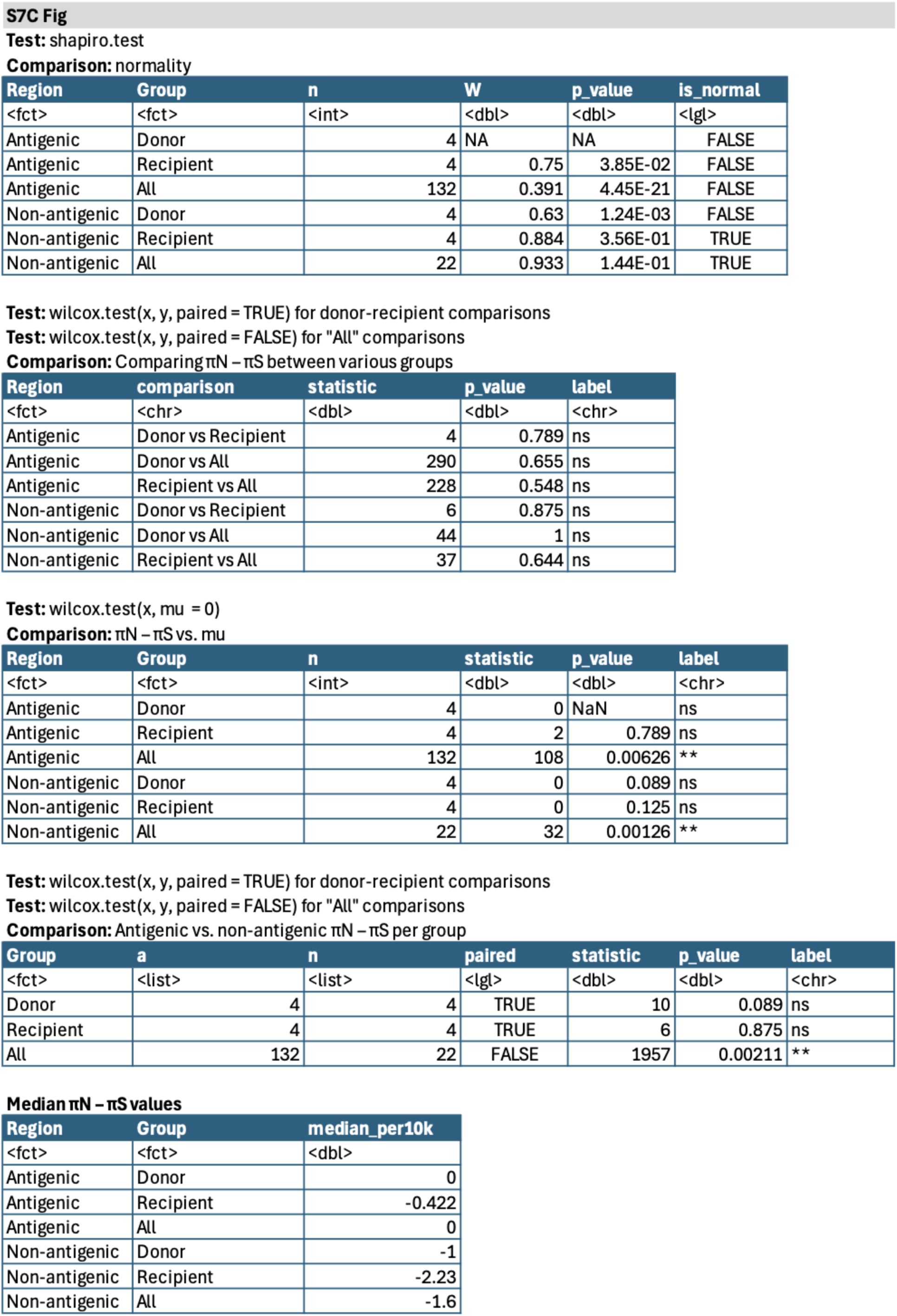
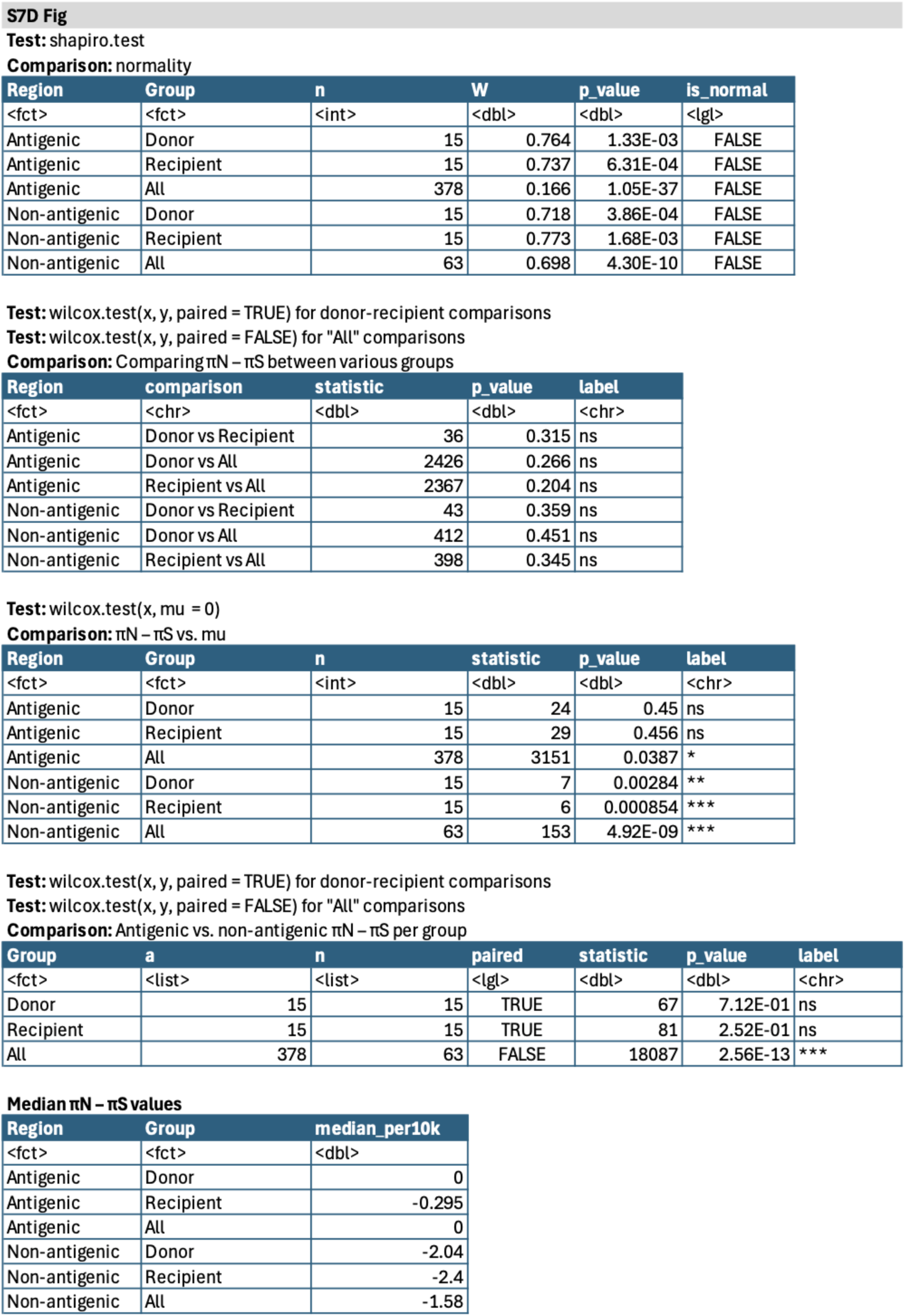
Statistical tests and results from the study. Genes shown: PB2 (polymerase basic 2), PB1 (polymerase basic 1), PA (polymerase acidic), HA (hemagglutinin), NP (nucleoprotein), and NA (neuraminidase). Not included: MP (matrix; M1/M2) and NS (nonstructural; NS1/NEP) due to overlapping reading frames.

**S4 Table.** Genomic positions for H3 antigenic sites. Genes shown: HA (hemagglutinin) only.

| Start | Stop | ID | Antigenic_Site | HA_number |
| --- | --- | --- | --- | --- |
| 429 | 431 | Antigenic | A | 122 |
| 435 | 437 | Antigenic | A | 124 |
| 441 | 443 | Antigenic | A | 126 |
| 453 | 455 | Antigenic | A | 130 |
| 456 | 458 | Antigenic | A | 131 |
| 459 | 461 | Antigenic | A | 132 |
| 462 | 464 | Antigenic | A | 133 |
| 468 | 470 | Antigenic | A | 135 |
| 474 | 476 | Antigenic | A | 137 |
| 477 | 479 | Antigenic | A | 138 |
| 483 | 485 | Antigenic | A | 140 |
| 489 | 491 | Antigenic | A | 142 |
| 492 | 494 | Antigenic | A | 143 |
| 495 | 497 | Antigenic | A | 144 |
| 498 | 500 | Antigenic | A | 145 |
| 501 | 503 | Antigenic | A | 146 |
| 513 | 515 | Antigenic | A | 150 |
| 519 | 521 | Antigenic | A | 152 |
| 567 | 569 | Antigenic | A | 168 |
| 447 | 449 | Antigenic | B | 128 |
| 450 | 452 | Antigenic | B | 129 |
| 528 | 530 | Antigenic | B | 155 |
| 531 | 533 | Antigenic | B | 156 |
| 534 | 536 | Antigenic | B | 157 |
| 537 | 539 | Antigenic | B | 158 |
| 540 | 542 | Antigenic | B | 159 |
| 543 | 545 | Antigenic | B | 160 |
| 552 | 554 | Antigenic | B | 163 |
| 555 | 557 | Antigenic | B | 164 |
| 558 | 560 | Antigenic | B | 165 |
| 621 | 623 | Antigenic | B | 186 |
| 624 | 626 | Antigenic | B | 187 |
| 627 | 629 | Antigenic | B | 188 |
| 630 | 632 | Antigenic | B | 189 |
| 633 | 635 | Antigenic | B | 190 |
| 639 | 641 | Antigenic | B | 192 |
| 642 | 644 | Antigenic | B | 193 |
| 645 | 647 | Antigenic | B | 194 |
| 651 | 653 | Antigenic | B | 196 |
| 654 | 656 | Antigenic | B | 197 |
| 657 | 659 | Antigenic | B | 198 |
| 195 | 197 | Antigenic | C | 44 |
| 198 | 200 | Antigenic | C | 45 |
| 201 | 203 | Antigenic | C | 46 |
| 204 | 206 | Antigenic | C | 47 |
| 207 | 209 | Antigenic | C | 48 |
| 213 | 215 | Antigenic | C | 50 |
| 216 | 218 | Antigenic | C | 51 |
| 222 | 224 | Antigenic | C | 53 |
| 225 | 227 | Antigenic | C | 54 |
| 882 | 884 | Antigenic | C | 273 |
| 888 | 890 | Antigenic | C | 275 |
| 891 | 893 | Antigenic | C | 276 |
| 897 | 899 | Antigenic | C | 278 |
| 900 | 902 | Antigenic | C | 279 |
| 903 | 905 | Antigenic | C | 280 |
| 945 | 947 | Antigenic | C | 294 |
| 954 | 956 | Antigenic | C | 297 |
| 960 | 962 | Antigenic | C | 299 |
| 963 | 965 | Antigenic | C | 300 |
| 975 | 977 | Antigenic | C | 304 |
| 978 | 980 | Antigenic | C | 305 |
| 984 | 986 | Antigenic | C | 307 |
| 987 | 989 | Antigenic | C | 308 |
| 990 | 992 | Antigenic | C | 309 |
| 993 | 995 | Antigenic | C | 310 |
| 996 | 998 | Antigenic | C | 311 |
| 999 | 1001 | Antigenic | C | 312 |
| 351 | 353 | Antigenic | D | 96 |
| 369 | 371 | Antigenic | D | 102 |
| 372 | 374 | Antigenic | D | 103 |
| 414 | 416 | Antigenic | D | 117 |
| 426 | 428 | Antigenic | D | 121 |
| 564 | 566 | Antigenic | D | 167 |
| 573 | 575 | Antigenic | D | 170 |
| 576 | 578 | Antigenic | D | 171 |
| 579 | 581 | Antigenic | D | 172 |
| 582 | 584 | Antigenic | D | 173 |
| 585 | 587 | Antigenic | D | 174 |
| 588 | 590 | Antigenic | D | 175 |
| 591 | 593 | Antigenic | D | 176 |
| 594 | 596 | Antigenic | D | 177 |
| 600 | 602 | Antigenic | D | 179 |
| 609 | 611 | Antigenic | D | 182 |
| 666 | 668 | Antigenic | D | 201 |
| 672 | 674 | Antigenic | D | 203 |
| 684 | 686 | Antigenic | D | 207 |
| 687 | 689 | Antigenic | D | 208 |
| 690 | 692 | Antigenic | D | 209 |
| 699 | 701 | Antigenic | D | 212 |
| 702 | 704 | Antigenic | D | 213 |
| 705 | 707 | Antigenic | D | 214 |
| 708 | 710 | Antigenic | D | 215 |
| 711 | 713 | Antigenic | D | 216 |
| 714 | 716 | Antigenic | D | 217 |
| 717 | 719 | Antigenic | D | 218 |
| 720 | 722 | Antigenic | D | 219 |
| 741 | 743 | Antigenic | D | 226 |
| 744 | 746 | Antigenic | D | 227 |
| 747 | 749 | Antigenic | D | 228 |
| 750 | 752 | Antigenic | D | 229 |
| 753 | 755 | Antigenic | D | 230 |
| 777 | 779 | Antigenic | D | 238 |
| 783 | 785 | Antigenic | D | 240 |
| 789 | 791 | Antigenic | D | 242 |
| 795 | 797 | Antigenic | D | 244 |
| 801 | 803 | Antigenic | D | 246 |
| 804 | 806 | Antigenic | D | 247 |
| 807 | 809 | Antigenic | D | 248 |
| 234 | 236 | Antigenic | E | 57 |
| 240 | 242 | Antigenic | E | 59 |
| 249 | 251 | Antigenic | E | 62 |
| 252 | 254 | Antigenic | E | 63 |
| 264 | 266 | Antigenic | E | 67 |
| 288 | 290 | Antigenic | E | 75 |
| 297 | 299 | Antigenic | E | 78 |
| 303 | 305 | Antigenic | E | 80 |
| 306 | 308 | Antigenic | E | 81 |
| 309 | 311 | Antigenic | E | 82 |
| 312 | 314 | Antigenic | E | 83 |
| 321 | 323 | Antigenic | E | 86 |
| 324 | 326 | Antigenic | E | 87 |
| 327 | 329 | Antigenic | E | 88 |
| 336 | 338 | Antigenic | E | 91 |
| 339 | 341 | Antigenic | E | 92 |
| 390 | 392 | Antigenic | E | 109 |
| 843 | 845 | Antigenic | E | 260 |
| 846 | 848 | Antigenic | E | 261 |
| 849 | 851 | Antigenic | E | 262 |
| 858 | 860 | Antigenic | E | 265 |

**S5 Table.** Genomic positions for H1 antigenic sites. Genes shown: HA (hemagglutinin) only.

| Start | Stop | ID | Antigenic_Site | HA_number |
| --- | --- | --- | --- | --- |
| 280 | 297 | Antigenic | Cb | 70 |
| 442 | 447 | Antigenic | Sa | 124 |
| 481 | 498 | Antigenic | Ca2 | 137 |
| 529 | 543 | Antigenic | Sa | 153 |
| 547 | 564 | Antigenic | Sa | 159 |
| 568 | 582 | Antigenic | Ca1 | 166 |
| 622 | 657 | Antigenic | Sb | 184 |
| 679 | 687 | Antigenic | Ca1 | 203 |
| 733 | 738 | Antigenic | Ca2 | 221 |
| 775 | 783 | Antigenic | Ca1 | 235 |

